# Releasing Protein Cargo from Synthetic Cells by Coupling Cell-Free Expression to Osmotic Actuation

**DOI:** 10.64898/2026.10.06.757017

**Authors:** Hongfei Liu, Jérémie J. M. Marlhens, Simon Schubotz, Heinz Koeppl

## Abstract

Macromolecular release from synthetic cells often relies on membrane pores or channels. We explored whether internally generated osmotic stress could increase membrane permeability and promote release. Using double-emulsion microfluidics, we encapsulated an *Escherichia coli* cell-free expression system in Pluronic L121 polymersomes to co-express the fluorescent protein mScarlet3 and the dextranase PsDex1711 in the presence of dextran. PsDex1711-mediated dextran hydrolysis produced a greater increase in bulk osmolality than reactions without PsDex1711 DNA. Tracking hundreds of individual polymersomes showed that PsDex1711-containing populations expanded, whereas control populations contracted. The distribution of normalized fluorescence revealed a subset with lower residual fluorescence than the main population. In selected expanded polymersomes, integrated internal mScarlet3 fluorescence declined to 18–22 % of peak levels, without visible membrane disruption, consistent with protein release. These findings support a proof of concept for using internal gene expression to promote macromolecular release through osmotic actuation without introducing dedicated membrane pores.

## Introduction

Synthetic cells combine cell-sized compartments with encapsulated biochemical systems to reconstruct selected cellular functions. ^1–3^ Their membranes retain biochemical machinery while mediating molecular exchange with the surroundings.^1,4–6^ By connecting internal reactions with the external environment, membrane transport supports sustained biochemical activity, biosensing, communication, and cargo release, making control over its selectivity and timing critical for developing functional synthetic cells. ^4,7–9^

Sustained cell-free protein synthesis has been supported by nutrient exchange through membrane-spanning *α*-hemolysin pores.^9^ Biosensing and inducible expression have been enabled by ion or inducer entry through osmotically activated MscL channels.^10,11^ Controlled cargo release has been achieved by combining cell-free pore expression with membrane-composition control^12^ or temperature-responsive genetic regulation.^13^ Therapeutic protein delivery has also been demonstrated using cell-free-expressed perfringolysin O pores. ^14^ In addition, sequential reactant delivery through interacting DNA-based pores has supported actin assembly and RNA transcription.^15^ These approaches require protein folding and assembly or DNA structure design, together with functional integration into membranes of compatible composition. ^15–20^

Osmotically driven membrane expansion offers another route to altering molecular transport. An increase in internal osmotic pressure relative to the surroundings can drive water influx and vesicle expansion. Internal reactions can generate this difference by increasing the number of solute molecules retained inside the vesicle.^21^ External hypotonic shock released internally expressed proteins from polymersomes through transient rupture and resealing,^22^ whereas internal photochemical cleavage induced osmotic rupture and content release.^21^ For synthetic cells, however, retaining vesicle structure and biochemical machinery during cargo release could preserve the capacity for further reactions. Molecular exchange can support sustained cell-free expression, ^9^ initiate expression through externally supplied components, ^23^ and enable sequential biochemical reactions.^15^ Cell-free expression also provides a means of regulating membrane transport through genetic control.^11,13^ We therefore asked whether osmotic changes generated by an internally expressed hydrolase could increase membrane permeability. Ideally, a membrane would withstand osmotic expansion without rupturing and have a porous structure with transport pathways that could widen upon stretching.

We selected Pluronic L121 (PL121), a PEO–PPO–PEO triblock copolymer forming porous membranes with low bending stiffness^24^ and intrinsic permeability to small hydrophilic solutes.^24,25^ Previous studies reported osmotic swelling and small-dye release, ^25^ while studies using PL121 polymersomes as compartments for enzymatic reactions reported an approximate membrane molecular-weight cutoff of 500 Da.^26,27^ This cutoff provides a guide for selecting reaction products that could accumulate inside polymersomes and generate a transmembrane osmotic pressure difference.

Accordingly, we produced PL121 polymersomes by double-emulsion microfluidics and encapsulated an *Escherichia coli* cell-free transcription–translation (TXTL) system, dextran 70 kDa (Dex70), and DNA encoding mScarlet3 and the dextranase PsDex1711 (Figure 1A). The fluorescent protein mScarlet3 served as both an expression reporter and cargo. PsDex1711 hydrolyzes Dex70 into isomaltooligosaccharides (IMOs; degree of polymerization 3–6, 504– 991 Da),^28^ near or above the reported cutoff^26,27^ (Figure 1B). Hydrolysis increases the number of solute molecules, whose accumulation could generate an osmotic pressure difference sufficient to drive water influx and polymersome expansion. We hypothesized that membrane stretching could then increase intrinsic permeability, allowing previously retained mScarlet3 to escape (Figure 1C). PsDex1711 expression significantly shifted the population response toward expansion, and selected expanding polymersomes showed a loss of integrated internal mScarlet3 fluorescence while retaining their overall morphology, consistent with protein release.

**Figure 1.**
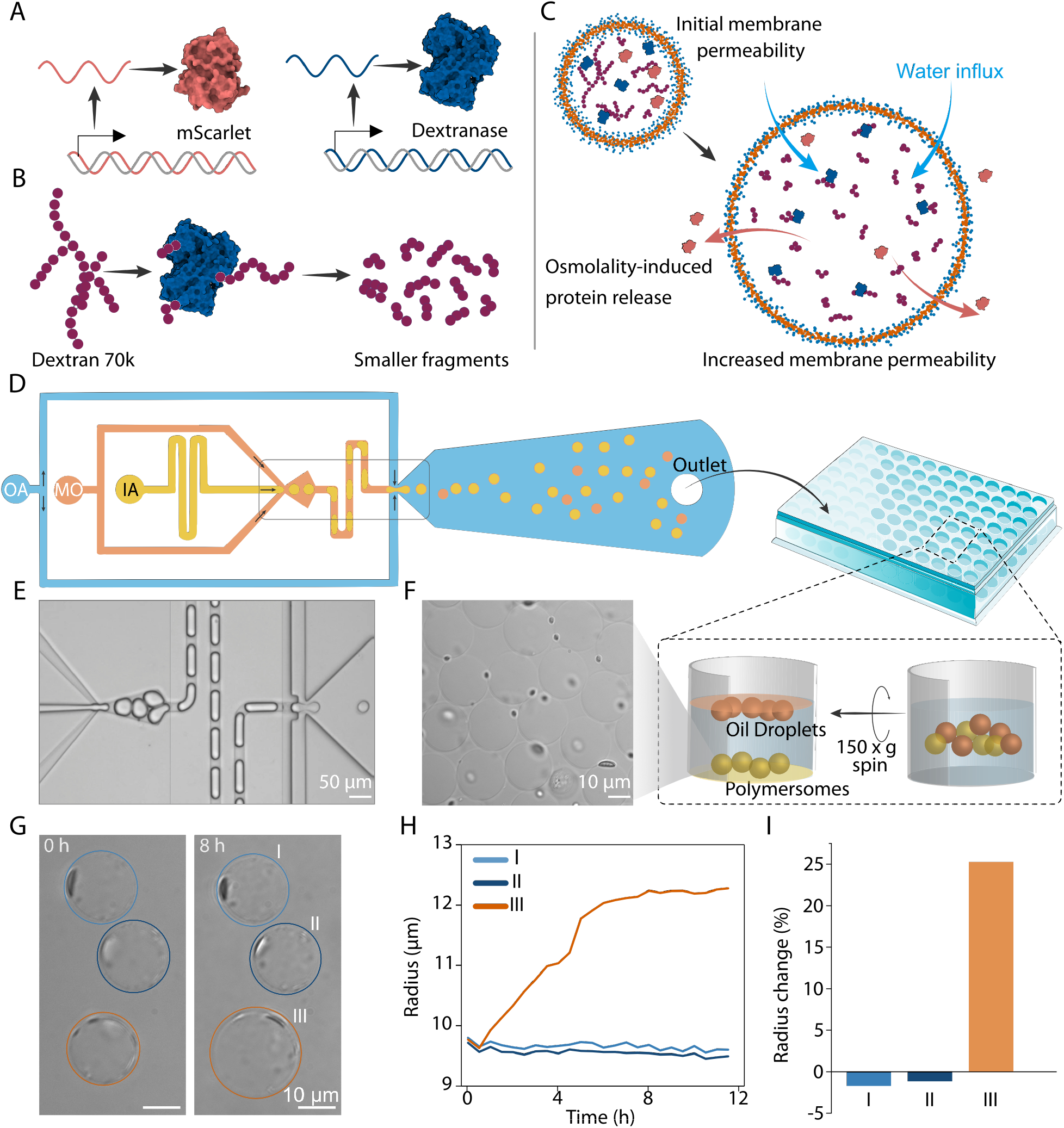
System design, polymersome preparation, and single-vesicle tracking. (A–C) Cell-free expression of the fluorescent protein mScarlet3 and the dextranase PsDex1711, Dex70 hydrolysis, and the proposed mechanism of osmotically driven protein release. (D) Microfluidic double-emulsion production, collection in 96-well plates, and centrifugation at 150 × *g* to separate the collected output into polymersomes near the well bottom and residual oil droplets near the surface. IA, inner aqueous phase; MO, middle oil phase; OA, outer aqueous phase. (E,F) Bright-field images of double-emulsion generation (E) and polymersomes after centrifugation (F). (G–I) Segmentation and tracking, absolute radius trajectories, and endpoint radius changes [(*R*_end_ − *R*_0_)*/R*_0_] ×100 % for three selected polymersomes from one recording.

## Results and Discussion

We produced PL121 polymersomes by microfluidic water-in-oil-in-water double-emulsion templating and collected them in 96-well plates containing the outer aqueous phase (Figure 1D,E). Centrifugation sedimented the denser polymersomes to the well bottom, while 1-octanol droplets generated as a byproduct during production remained near the surface. This separation enabled imaging through the plate bottom without visible oil droplets (Figure 1D,F).

Three selected polymersomes from one recording illustrate single-vesicle segmentation and tracking (Figure 1G), the resulting radius trajectories (Figure 1H), and endpoint radius changes (Figure 1I).

We first tested whether PL121 polymersomes prepared using 1-octanol retained macro-molecules, as reported for those prepared with other solvents.^25–27^ FITC–mPEG 2 kDa retention was assessed in PBS-based buffer without the TXTL system to minimize osmotic changes and membrane deformation that could alter membrane permeability. The observed polymersome showed no apparent size change and retained 88 % of its initial integrated fluorescence after 12 h (Figure 2A,B). This supports substantial macromolecular retention. Photobleaching may have contributed to the modest fluorescence decline (Figure S12).

**Figure 2.**
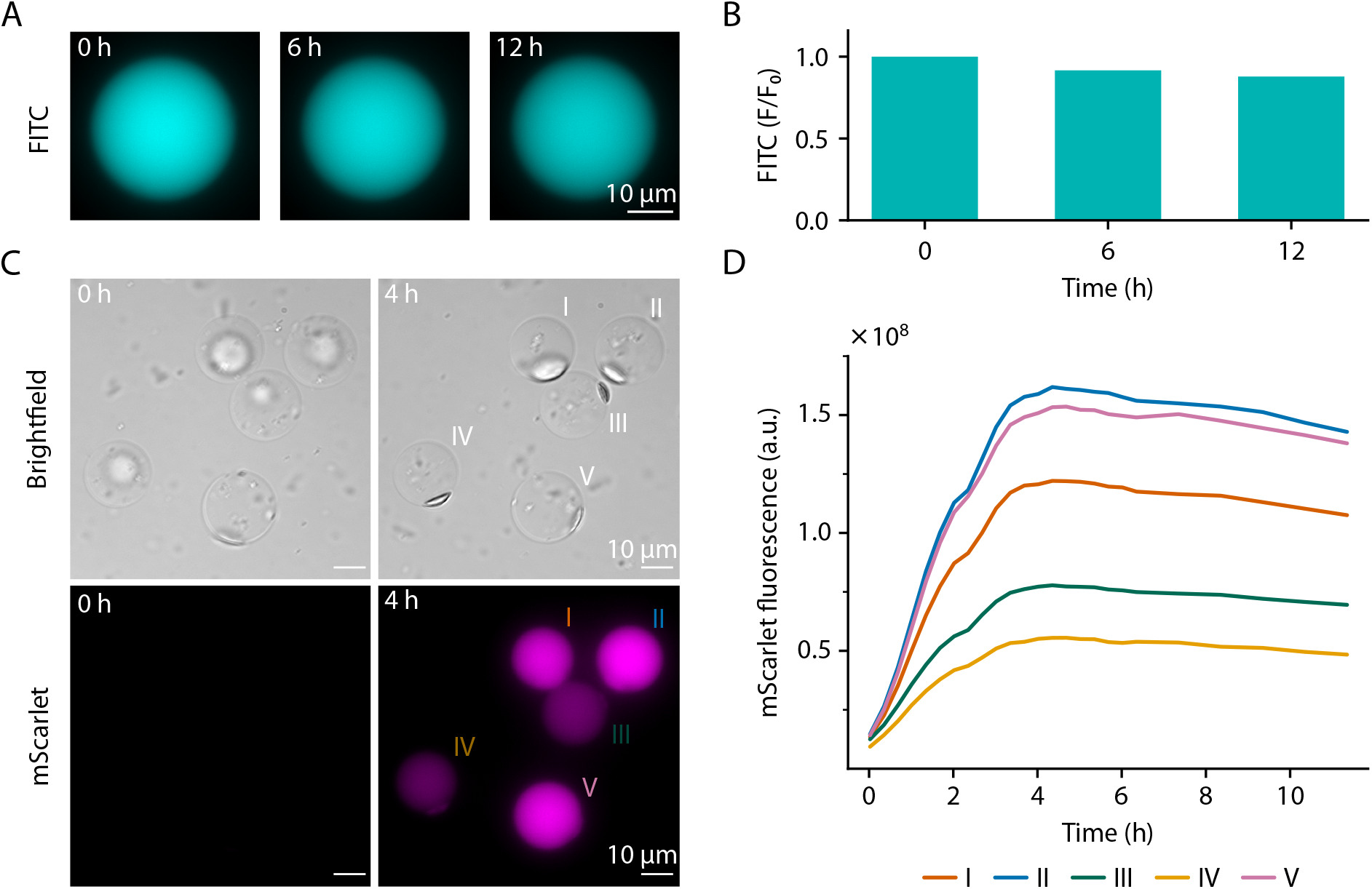
Macromolecular retention and cell-free expression in PL121 polymersomes. (A) Fluorescence images of one representative polymersome containing FITC–mPEG 2 kDa at 0 h, 6 h and 12 h in a tracer-retention assay without the TXTL system. (B) Integrated fluorescence of the polymersome in A, normalized to its initial value (*F/F*_0_). (C) Bright-field (top) and mScarlet3 fluorescence (bottom) images at 0 h and 4 h of polymersomes containing mScarlet3 DNA and Dex70 without PsDex1711 DNA. (D) Integrated mScarlet3 fluorescence trajectories corresponding to the five labeled polymersomes in C.

In polymersomes containing *E. coli* TXTL, mScarlet3 DNA, and Dex70 without PsDex1711 DNA, the development of lumenal mScarlet3 fluorescence demonstrated cell-free protein expression (Figure 2C). Integrated fluorescence reached a peak after ∼4 h and subsequently declined by 10–13 % over the remaining observation period (Figure 2D). Despite similar initial radii, with the largest only 10 % greater than the smallest, and similar fluorescence time courses, the five tracked polymersomes differed by 2.9-fold in peak integrated fluorescence, indicating heterogeneity in expression.

We examined whether cell-free PsDex1711 expression increased osmolality in Dex70-containing reactions. Bulk osmolality increased relative to initial values in all three conditions at 8 h and overnight. Mean osmolality increases with PsDex1711 were approximately 1.7 and 1.6 times those in the mScarlet3-only control at 8 h and overnight, respectively, reaching 115 mOsm kg^−1^ overnight (Figure 3A).

**Figure 3.**
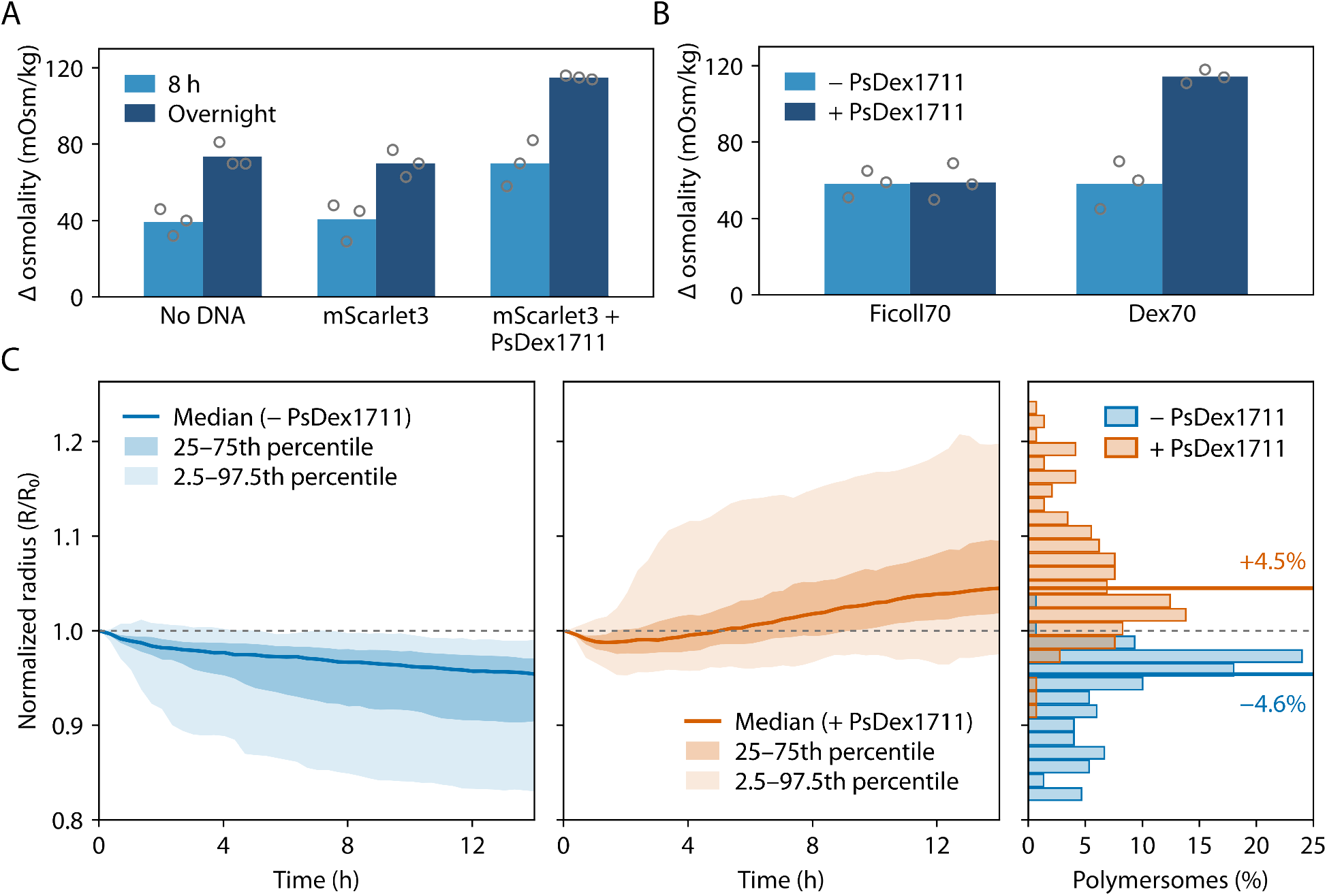
Bulk osmolality changes and polymersome size responses associated with cell-free PsDex1711 expression. (A) Osmolality changes after 8 h and after overnight incubation in Dex70-containing cell-free reactions with no added DNA, 5 nM mScarlet3 DNA alone, or 5 nM mScarlet3 DNA plus 30 nM PsDex1711 DNA. Mean increases with PsDex1711 were approximately 1.7 and 1.6 times those in the mScarlet3-only control at 8 h and overnight, respectively. (B) Overnight osmolality changes in Ficoll70- or Dex70-containing cell-free reactions with 5 nM mScarlet3 DNA, in the absence (−PsDex1711) or presence (+PsDex1711) of 20 nM PsDex1711 DNA. PsDex1711 expression produced an additional increase with Dex70 but not with Ficoll70. In A and B, Δ osmolality denotes the change relative to each reaction’s initial value. Columns show means, and open circles show individual values from three independent experiments (*n* = 3). (C) Normalized radius trajectories (*R/R*_0_) of mScarlet3-only controls (−PsDex1711; left; *n* = 150 polymersomes) and PsDex1711-containing polymersomes (+PsDex1711; middle; *n* = 145 polymersomes), and their distributions at 14 h (right). Solid curves show medians; darker and lighter shading indicate the 25th–75th and 2.5th–97.5th percentile ranges, respectively. Horizontal colored lines in the right plot mark endpoint medians, showing contraction in controls and expansion with PsDex1711. Percentages indicate median radius changes relative to the initial value.

To determine whether this additional osmolality increase depended on Dex70 hydrolysis rather than PsDex1711 expression alone, we replaced Dex70 with Ficoll70, a polysaccharide of similar nominal molecular weight that is not a PsDex1711 substrate. In Ficoll70-containing reactions, overnight osmolality increases were similar regardless of PsDex1711 DNA addition. Both increases were close to that in the Dex70 group without PsDex1711 DNA. Only the combination of PsDex1711 expression and the substrate Dex70 resulted in a larger increase, approximately twice that in the Dex70 control (Figure 3B). Together, these experiments support PsDex1711-mediated Dex70 hydrolysis as the source of the additional osmolality increase.

We next examined whether these reactions could drive polymersome expansion. Before polymersome production, the outer-phase osmolality was 20–30 mOsm kg^−1^ higher than that of the inner phase. Both populations initially contracted, consistent with this initial imbalance. The mScarlet3-only controls continued contracting, whereas the PsDex1711-containing population recovered and its median normalized radius rose above unity (Figure 3C). At 14 h, median radius changes were −4.6 % in controls and 4.5 % in PsDex1711-containing polymersomes. The distributions differed significantly between the two populations in this experiment (two-sided Mann–Whitney U test, *p <* 0.001). Radius changes were predominantly negative in controls and positive with PsDex1711, indicating a population-level shift from contraction toward expansion (Figure 3C). Additional independent experiments are presented in Figure S5 and Table S2.

Continued control contraction despite rising bulk osmolality highlights why bulk changes cannot be directly assigned to the lumen or used to predict expansion. Encapsulated reactions can differ in expression output (Figure 2D) and reaction progress, while outward permeation limits solute accumulation inside polymersomes and thus the buildup of a transmembrane osmotic pressure difference. TXTL reactions without PsDex1711 may generate small solutes that raise bulk osmolality but readily cross the membrane, limiting their contribution to a sustained transmembrane osmotic difference. Larger oligosaccharides from Dex70 hydrolysis could be retained long enough to drive water influx. We adjusted outer-phase osmolality with dextran 1 kDa (Dex1k), expected to provide a more persistent external osmotic contribution than readily permeable solutes; the initial 20–30 mOsm kg^−1^ offset was chosen empirically to keep controls contracting.

We next investigated whether osmotic expansion could enable release of internally expressed mScarlet3. We quantified integrated fluorescence rather than mean intensity, which can decrease through dilution as a polymersome expands. The distribution of endpoint fluorescence normalized to each polymersome’s peak showed a small subset with markedly lower values, separated from the main population by a gap (Figure S6). To investigate whether this pronounced fluorescence loss reflected release, we examined individual fluorescence and size trajectories together with the corresponding images. In the representative mScarlet3-only DNA control, lumenal fluorescence developed and remained bright, while the polymersome gradually contracted by ∼10 % in radius (Figure 4A,C,D).

**Figure 4.**
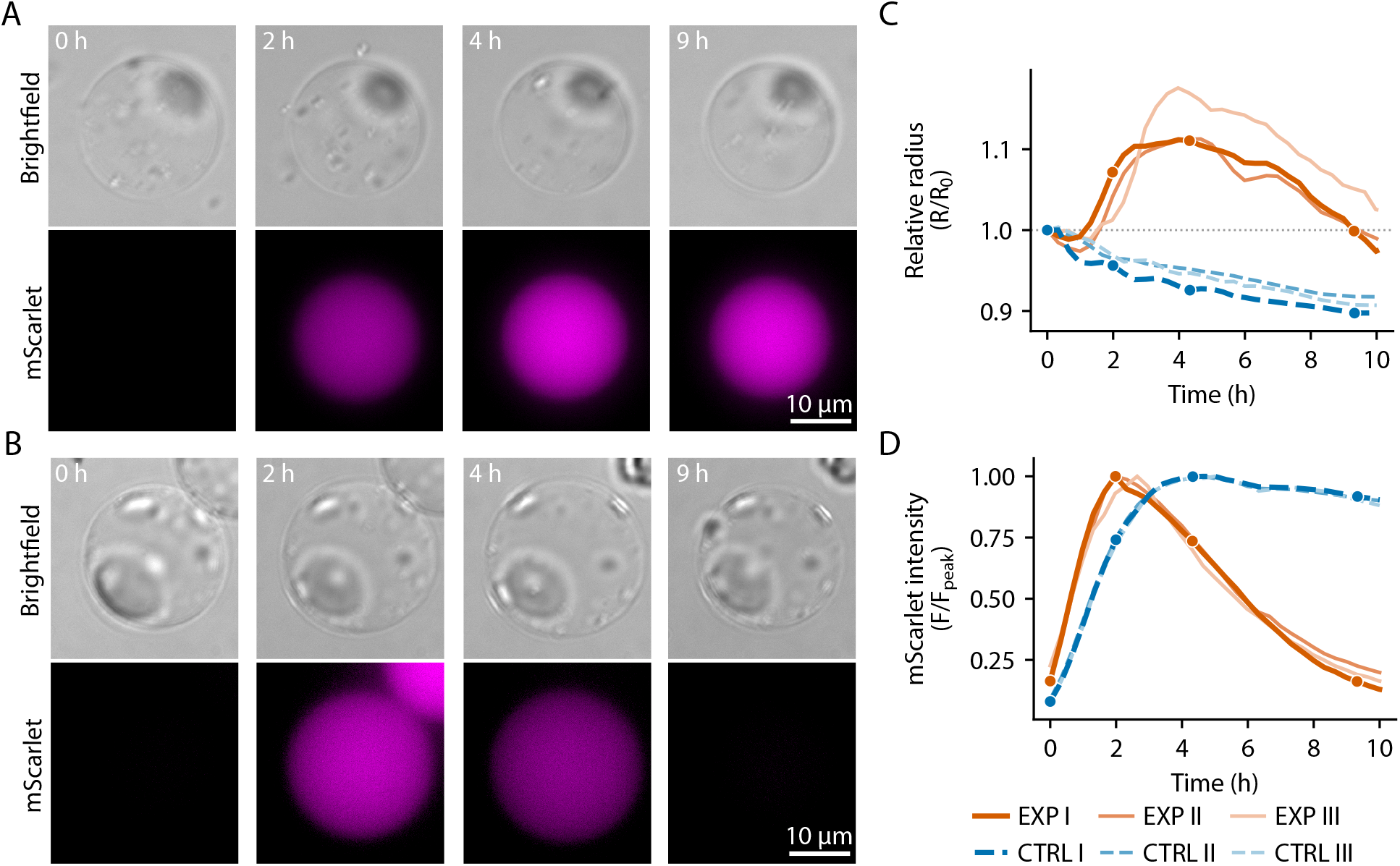
Polymersome size dynamics and loss of internally expressed mScarlet3. (A,B) Bright-field (top) and mScarlet3 fluorescence (bottom) images at 0 h, 2 h, 4 h and 9 h of a representative mScarlet3-only control (A; CTRL I) and a PsDex1711-containing polymersome (B; EXP I). Image times illustrate the initial state, fluorescence maximum, maximum radius, and subsequent fluorescence decline of EXP I, with the nearest available times shown for CTRL I. (C,D) Radius normalized to its initial value (*R/R*_0_; C) and background-corrected integrated mScarlet3 fluorescence normalized to its maximum over the full recorded trajectory (*F/F*_peak_; D) for three selected polymersomes per condition. Orange solid lines indicate PsDex1711-containing polymersomes; blue dashed lines indicate mScarlet3-only controls. Thick lines correspond to EXP I and CTRL I; markers indicate the image times in A and B. Selected PsDex1711-containing polymersomes showed transient expansion followed by pronounced fluorescence loss, whereas controls contracted and retained high fluorescence after its initial accumulation.

In contrast, the representative PsDex1711-containing polymersome expanded while lumenal mScarlet3 fluorescence developed and reached its peak at ∼2 h, followed by a pronounced fluorescence decline (Figure 4B). At approximately 4 h, the radius reached its maximum, 11 % larger than its initial value, while fluorescence had decreased to 78 % of its peak (Figure 4C,D). Continued expansion during fluorescence decline suggests that membrane permeability increased while the compartment remained intact, with ongoing dextran hydrolysis potentially sustaining the osmotic driving force. By approximately 9 h, fluorescence had declined to 18 % of its peak, close to its initial level, with the radius also near its starting value and no visible membrane defects in bright-field images. Two additional selected PsDex1711-containing polymersomes showed similar expansion and fluorescence-loss trajectories. Across these three polymersomes, maximum radius increases were 11–18 %, and fluorescence subsequently declined to 18–22 % of peak, compared with 90–92 % in three controls (Figure 4C,D; images in Figures S7–S10). Together, these observations suggest that osmotic expansion increased membrane permeability, allowing release of internally expressed mScarlet3 without apparent membrane rupture.

However, some polymersomes expanded substantially while retaining strong mScarlet3 fluorescence, indicating that expansion did not consistently lead to observable release (Figure S11).

Previous work on PL121 polymersomes reported a polymer reservoir in the membrane region previously occupied by an oil pocket that disappeared during swelling. ^25^ Recruitment of this material into the membrane could accommodate expansion with less membrane stretching, potentially limiting the increase in permeability.

Expression heterogeneity was observed among polymersomes (Figure 2D), consistent with reports in other compartmentalized cell-free systems. ^29,30^ Although PsDex1711 expression was not measured in individual polymersomes, differences in enzyme expression could affect the rate and extent of Dex70 hydrolysis. Hydrolysis inside polymersomes occurs alongside solute exchange across the membrane, changing the transmembrane osmotic difference over time. Differences in these coupled processes, enzyme expression, and available polymer reservoirs could limit the fraction of polymersomes reaching conditions sufficient for protein release.

In summary, this study provides a proof of concept for using internal gene expression to drive osmotic actuation and promote macromolecular release in PL121 synthetic cells without introducing dedicated membrane pores. PsDex1711 expression shifted the population response from contraction to expansion. Within this population, a subset showed greater fluorescence loss, and analysis of selected individuals supported a mechanism of protein release through increased membrane permeability while maintaining overall structural integrity. Incorporating inducible expression and feedback regulation could enable more complex control of synthetic-cell signaling and the release of functional macromolecules.

## Methods

### Cell-Free Reactions

*E. coli* TXTL reactions were prepared using laboratory-made crude extract and energy buffer according to a protocol adapted from Sun et al.^31^ Unless otherwise stated, encapsulation reactions contained crude extract (33 % v/v), energy buffer (33 % v/v), linear mScarlet3 DNA (5 nM), Dex70 (70 mg mL^−1^), and nuclease-free water. Dextranase-containing reactions additionally contained linear DNA encoding the PsDex1711-derived construct (30 nM).^28^ FITC–mPEG 20 kDa (10 µM) was included in tracer experiments. Reactions were encapsulated without pre-expression.

Bulk mScarlet3 fluorescence was monitored at 28 ^°^C using a CLARIOstar Plus microplate reader (BMG LABTECH). For each PsDex1711 polymersome expansion experiment, bulk osmolality of the corresponding inner-phase formulation was measured in separate aliquots immediately after preparation and after overnight incubation at 28 ^°^C using a freezing-point osmometer (Osmomat 3000 Basic, Gonotec). Complete formulations, reagent sources, DNA sequences, and protocol modifications are provided in the SI.

### PL121 Polymersome Preparation

PL121 polymersomes were produced by double-emulsion templating in a custom poly(dimethylsiloxane) (PDMS) microfluidic device based on established methods.^32–35^ The inner phase contained the TXTL reaction described above, the middle phase contained PL121 and 1-octanol (3:97 v/v), and the outer phase contained poly(vinyl alcohol) (PVA; 2 % w/v) and energy buffer (33 % v/v). Before production, inner-phase osmolality was typically 550–600 mOsm kg^−1^. The outer phase was adjusted with Dex1k and verified to be 20– 30 mOsm kg^−1^ higher. Tracer-only assays used FITC–mPEG 2 kDa in PBS-based inner and outer phases without the TXTL system. Device fabrication, reagent sources, and complete phase compositions are provided in the SI.

### Microfluidic Operation and Imaging

The three phases were independently supplied using a LineUp pressure-control system (Fluigent). Initial pressures were typically 650 mbar, 390 mbar and 420 mbar, respectively, and adjusted for stable double-emulsion production. Samples were collected into 96-well plates containing 100 µL of outer phase and centrifuged twice at 150 × *g* for 3 min each (Megafuge ST4R Plus, Thermo Fisher Scientific), overlaid with anti-evaporation oil, and sealed. Imaging used an inverted microscope (Ti, Nikon) with a SPECTRA X light engine (6-LCR-SA, Lumencor) and an sCMOS camera (ORCA-Fusion C14440-20UP, Hamamatsu Photonics). Overview imaging used a 10×/0.45 NA objective; time-lapse imaging used a 60×/1.40 NA oil-immersion objective. Time-lapse imaging was performed at 28 *±* 2 ^°^C every 20 min, typically for 14 h and up to 18 h. GFP- and mCherry-channel exposures were 50 ms and 200 ms, respectively.

### Image Analysis

An automated single-polymersome analysis pipeline was implemented within an Electronic Laboratory Notebook (ELN) framework. ^36^ A custom Cellpose model was trained on combined enhanced bright-field and normalized fluorescence channels to improve membrane detection.^37^ Masks were filtered by size, circularity, solidity, and eccentricity to exclude artifacts and irregular objects. The perimeter-equivalent radius was calculated from each valid mask as *R*_eq_ = *P/*(2*π*), where *P* is the perimeter. Trajectories were reconstructed using trackpy (0.7), linking masks across frames despite changes in segmentation labels.

Integrated mScarlet3 and FITC–mPEG fluorescence was quantified within each valid mask in raw images after subtraction of an image-wide background baseline at each frame. Polymersome paths, radius trajectories, and fluorescence traces were inspected for each video in the ELN-integrated Plotly dashboard, with closer inspection of selected cases in Fiji/ImageJ.^38^ Segmentation thresholds, tracking parameters, background correction, and software versions are detailed in SI Section S3; analysis code and processed data is available in the accompanying repository.^39,40^

## Associated Content

### Supporting Information

The Supporting Information is available free of charge at the ACS Publications website.

Detailed TXTL preparation; DNA template sequences and amplification; microfluidic device fabrication and dimensions; phase formulations and polymersome preparation; image presentation, segmentation, tracking, and fluorescence quantification; dextranase construct and DNA-concentration screening; additional bulk expression and osmolality data; population size responses and statistics across independent experiments; peak-normalized fluorescence distributions; supplementary single-polymersome images and trajectories; FITC–mPEG fluorescence under different imaging schedules; and reagent sources (PDF).

## Author Information

### Corresponding Author

Heinz Koeppl, Self-Organizing Systems Laboratory, Department of Electrical Engineering and Information Technology, Technische Universität Darmstadt, Merckstrasse 25, 64283 Darmstadt, Germany.

### Author Contributions

H.L. conceived the study, established the microfluidic platform, designed and performed the main experiments, acquired the experimental data, optimized polymersome preparation, cell-free encapsulation, and time-lapse imaging workflows, processed and analyzed data, prepared figures, and wrote the initial manuscript draft and coordinated manuscript finalization.

J.J.M.M. contributed to experimental design and cell-free experimental development, provided the cell-free expression system, and designed and constructed DNA templates with input from H.L. He performed DNA amplification, screening, and energy-buffer optimization with H.L. and contributed to data analysis, figure preparation, and manuscript development.

S.S. advised on time-lapse imaging, adapted the electronic laboratory notebook platform he had previously built, developed image segmentation, tracking, and data-processing workflows, processed and analyzed data, prepared figures, and contributed to manuscript development, particularly its structure and scientific argumentation.

H.K. provided conceptual guidance, supervised the project, and supported the research through funding and infrastructure.

All authors revised and approved the final manuscript.

### Notes

The authors declare no competing financial interest.

## Acknowledgments

J.J.M.M. was supported by the MSCA-DN SYNSENSO (grant agreement no. 101072980). We thank Ali Manshadi for technical assistance with CAD modifications and microfluidic chip fabrication, optimization of silicon wafer master fabrication parameters, and provision of fabrication details for the Supporting Information. We thank Bin Ji for helpful discussions and feedback on figure presentation. We thank İrem Ergenlioğlu for helping with the cell-free preparation and proofreading the manuscript. ChatGPT (OpenAI) and Gemini (Google) were used to assist with English-language revision based on author-provided content, manuscript checking and formatting, and Python code development for data analysis and plotting. The authors determined the scientific content and analytical approaches, reviewed and revised the AI-assisted text and code, and take responsibility for the manuscript and reported results.

## TOC Graphic

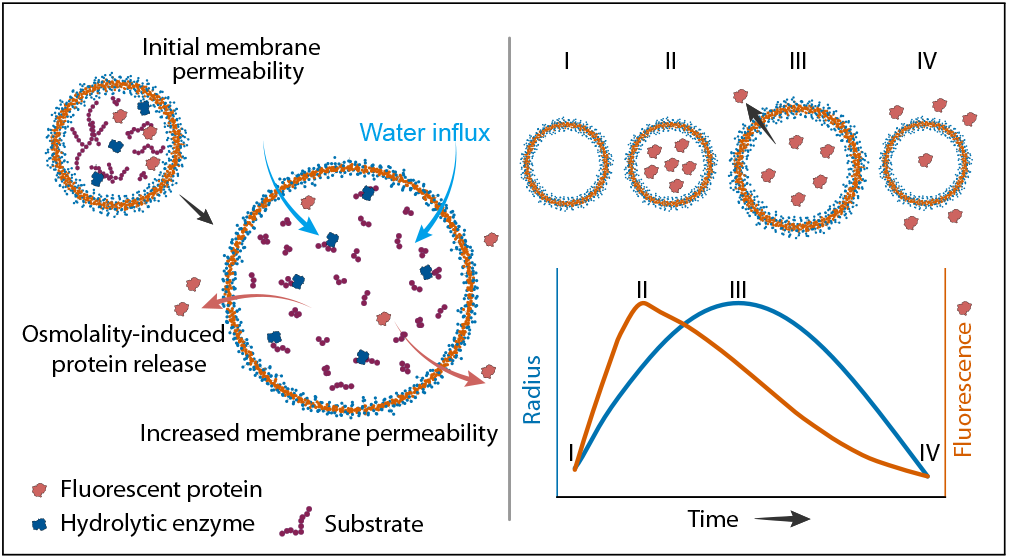

## Notes

https://doi.org/10.5281/zenodo.23185690

### Competing Interest Statement

The authors have declared no competing interest.

