## Supporting Information for "Releasing Protein Cargo from Synthetic Cells by Coupling Cell-Free Expression to Osmotic Actuation"

#### S1 DNA templates and cell-free assays

##### S1.1 Linear DNA templates

Four linear DNA templates were used in this study: one encoding mScarlet3 and three encoding dextranases evaluated in the bulk osmolality screen. All templates were constitutively expressed from the *BBa\_J23119* promoter and terminated by T500, and were used as linear DNA templates in cell-free reactions without a plasmid backbone.

The mScarlet3 template encoded mScarlet3 as the expression reporter and fluorescent protein cargo and was assembled by polymerase cycling assembly from individual parts. Three dextranase templates were evaluated: a dextranase from *Arthrobacter* sp. CB-8,<sup>1</sup> PsDex1711 from *Pseudarthrobacter* sp. RN22,<sup>2</sup> and CeDex2365-49 from *Cellulosimicrobium* sp. THN1.<sup>3</sup> The dextranase coding sequences were codon-optimized for *E. coli* K-12. For the *Arthrobacter* sp. CB-8 dextranase and PsDex1711, N-terminal signal peptides predicted using SignalP 6.0 were removed prior to codon optimization and synthesis.<sup>4</sup> For CeDex2365, the previously reported active GH49 fragment CeDex2365-49 (A24–A637) was used.<sup>3</sup> The dextranase templates contain the original UTR1 sequence, whereas the mScarlet3 template contains a single-nucleotide UTR1 variant, designated UTR1b here. PsDex1711 was selected for subsequent polymersome experiments based on its performance in the bulk osmolality screen (Figure S3).

All four templates contained the same 22-bp Twist 5' and 3' adapter sequences used as PCR primer-binding sites. These sites are highlighted separately in Table S1, which gives the full DNA sequences and constituent genetic parts.

##### S1.2 Linear DNA amplification

All linear DNA templates were amplified by PCR using repliQa HiFi ToughMix (Quantabio) with primers targeting the flanking Twist adapter sequences: forward, 5'-CAATCCGCCCTCACTACAACCG-3', and reverse, 5'-TCCCTCATCGACGCCAGAGTAG-3'. PCR products were purified using the NucleoSpin Gel and PCR Clean-up kit (Macherey–Nagel) and used directly as linear DNA templates for cell-free reactions.

**Table S1.** Full sequences of the linear DNA templates used in this study. Sequences are given 5'→3' in lowercase. The 22-bp Twist adapter sequences used as PCR primer-binding sites are highlighted separately; the 3' adapter shown in the forward-strand sequence is the reverse complement of the reverse primer. Color code: promoter, RBS/5' UTR, CDS, terminator, PCR primer-binding site; uncolored bases are other flanking/linker sequences. Coding sequences include the start and stop codons. UTR1 corresponds to the T7 gene 10 leader sequence used in the Noireaux TXTL system,<sup>5</sup> whereas UTR1b denotes the single-nucleotide variant present in the mScarlet3 template, containing an A-to-G substitution immediately downstream of the Shine–Dalgarno sequence. The T500 copies in the dextranase templates carry a shortened poly(T) tract introduced by the synthesis provider. Parts are from the Registry of Standard Biological Parts<sup>6</sup> (BBa\_J23119), Shin and Noireaux<sup>5</sup> (UTR1, T500), Gadella et al.<sup>7</sup> (mScarlet3), Okushima et al.<sup>1</sup> (*Arthrobacter* sp. CB-8 dextranase), Wang et al.<sup>2</sup> (PsDex1711), and Xu et al.<sup>3</sup> (CeDex2365-49).

| Construct | Sequence (5' → 3') | bp |
| --- | --- | --- |
| mScarlet3<br>J23119–UTR1b–<br>mScarlet3–T500 | caatccgccctcactacaaccgttgacagctagctcagtcctaggtataatgctagcaactg<br>agacagaggcacaataattttgtttaactttaagaaggaggtataccatggattccaccgaa<br>gccgtgattaaagagttcatgcgttttaaagtgcacatggaggggtcaatgaacggccatga<br>atttgagatcgaaggtgaaggagaaggccgcccgtacgagggcacccagaccgccaaactgc<br>gtgtgactaaaggtggccccctgccattttcgtgggatatcctgagcccgcagtttatgtac<br>ggtagccgggctttcactaagcatccggccgatattcccgattactggaagcaaagctttcc<br>ggaaggcttcaaatgggagcgtgtcatgaacttcgaagatggaggcgcggtcagcgttgcgc<br>aagatacaagtctcgaagatgggacgcttatttataaagttaaactgcgaggtacaaacttt<br>ccgccggacggccctgttatgcagaagaaaacgatggggtgggaagcgtctacggagcgcct<br>gtatcctgaagatgtggtactaaagggcgacattaaaatggcactgcgcttaaaagacggag<br>gccgctatttggcagatttcaaaaccacctatcgtgcgaagaaaccagtgcagatgccgggt<br>gcctttaatattgaccgcaaattagacatcaccagccataatgaagattatacggtggtcga<br>acagtacgaacgcagtgtagcgcgtcacagcaccggcggttcaggtggctcctaaacatcga<br>agatgtcggtagaagcccaaagcccgccgaaaggcgggcttttttttctactctggcgtcg<br>atgaggga | 876 |

Table S1 continued

| Construct | Sequence (5' → 3') | bp |
| --- | --- | --- |
| *Arthrobacter* sp.<br>CB-8 dextranase<br>J23119-UTR1-dextranase-T500 | caatccgccctcactacaaccgggtctctctatttgacagctagctcagtcctaggtataat<br>gctagcgctagcaataattttgtttaactttaagaaggagatataccatggccactgctccg<br>gcggcggcgcctccgggcgttccagctgcgttgaaggctgagcgggccattaccaccgtgga<br>caatggtaacctgcatacatggtggcacgataatggcgttttctcaccagcgacgcctacgc<br>agtcgtccgaagtgcgccgcagcagcttctacgacgtccaggttgcccaagcaaatcagccg<br>cagaaactgtatgatgccttcagctatatgagtatcccgcgttcgggaaaaggcaagattgg<br>ctatacggaagaagacggagctgaattttctagcgatgcccgcctaaccatgagttggtcct<br>cttttgagtatgcgaaggatgtgtgggttgaagtgagtttgcggaccggtcaaacaattagc<br>agtgcagatcaggttcagattcgaccgtcatcgtacaatttcgaaaaacagctggttgacgc<br>cgatacggtccgtatcaaagttccgtattcggatgccggctaccgctttagcgtcgaattcg<br>agccgcaactctacacagcctacaatgatatgtctggcgattcaggcaaattaaccaccgag<br>gccgctggcaatcgcccaatccataccgagccgcgcaattccatgatggtgttcgcggaacc<br>caaactgcgtggagaacaaaaagaacgtttagttcccacggaggaatcgggatcaatccact<br>atcccgaaccgggggaagtgcgcaatctgaatagtgtatccgaagagattatttactttcgt<br>ccaggaacgtattccatgggtccggactaccatgcagtactgcccgccaacgtaaagtgggt<br>gtatctggcgccgggcgcgtacgtcaagggcgccttccgtttcctccacgatacacagagcc<br>aatataaggtgactgggtacggcgtgcttagcggtgaacagtacgtctacgaagcggatacc<br>aataactcttatcatcatctgagcggcgcaagcaactgtcatagctcatgtgtaaaaatgtt<br>acaatttgcgtcagccgatgccgagcaaaaattagacttacagggcgtgaccgtagcagagc<br>ctccgtatcattcgtttgtggtctatggcaacgaacagacgtttcacatgaatgtggagaat<br>tataaacaggtgggcagctggtattggcagaccgatggtattgaactgtacaaaggttcaac<br>catgaaaaacaccttctttaacgcgaatgatgatgtgcttaaaatgtaccacagcgacgtca<br>ccatcgacaataccgtgatttggaaaaatgaaaacggtccggtgattcagtgggggtggacc<br>ccgcgtaacatagacaacgtgaacgttgcgaacacgacggttattcacaatcgcatgtattg<br>gaaagatgtaaaatataatacctgcatttttaatagcagtagccattgggaggacatgggca<br>gcacaacaaaagctgatccgaacaccactgtgaagaacatgcgctttgaaaacacggcagta<br>gaaggtatgactaactgcgcaatacgcgtttacgccctgtcggatacggaaaacattcatat<br>taaaaactttaacattggtgcctggaatggcctggaatggacctctcaggtgagccacctca<br>aaaggtatactaacagtgccggcgaaaaagtgaccatcggaaacgaagttcctgatgggaac<br>gggttggcactggagaattactcggtcggtggccaagtaatcgaaaagactggtgggaactc<br>ctcggattaccagctgggccgtcttggttttgacggcgaaaattgggaaaactggaatgcgt<br>ggaaatctgcgccgtaactcgagcaaagcccgccgaaaggcgggcttttctgttgtttgaga<br>ccctactctggcgtcgatgaggga | 2008 |

Table S1 continued

| Construct | Sequence (5' → 3') | bp |
| --- | --- | --- |
| PsDex1711<br>J23119-UTR1-PsDex1711-T500 | caatccgccctcactacaaccgggtctctctatttgacagctagctcagtcctaggtataat<br>gctagcgctagcaataattttgtttaactttaagaaggagatataccatgacggaacaaacg<br>ggctttactgcgtcaggtcctgaactacgtacctggtggcataacaattacgaatacaatgc<br>gaccagcccgacccagaatggcaatgttcgtcgcagttcattttacgaggttcaagtggcaa<br>ccgcggcggcaccggaaactcgctatgattcctttgcgtatatgagcatcccgcgttctgga<br>aagggcaaaataggttacacagaacctgatggggctgagttcagcagctcagccaatctctc<br>aatgtcgtggtcgagcttcgaatatagcaccgctgtgtgggtggatgtgaagttaacaactg<br>gccagaccatcacctcagttgaccaggtgagcattaggccaagtaaatatacctttgaaaaa<br>cacttggtggatgcgaatacggtccgtattaaagtgccatattcattatctggctaccgctt<br>cagcgttgagtttgaaccgcagaactttacggcctacaacgatatgtctggtactgcaggcg<br>tcctaacagaaaccggtggtggcgatcatcgcgcgatccatactgagccgcgcaacagcatg<br>atgatctttgcagagccgacgctgggaggaggagaggcagatcgtctgattcccacgagtgc<br>tagcggtaccattaactatccgcaggaagggctggtggataacctgggcagcgttacggaag<br>aggttatttatttccgtccgggtacctattacatgggttccggccaccgagcgagtatgccc<br>ccaaacgtaaaatggatctatctggcgccaggcgcctacgtcaaaggtgccttttacttccc<br>gagctccacgcagggcgtatataaagtgaccgggcgcggtgttctgagtggggaacagtacg<br>tgtatgaagcagatacgacaacgtctggatatccgcatagtactggcgccaattgtcacaat<br>acctgtgtaaagatgctggaattctcttcctcgtcaaccatgcagcaatatctggacgtcga<br>agggattaccatttctgaaccgccttaccatagctttgtgatttacgggccgcagaatgctt<br>atgacatggagatgcgcgtggacaactataaacaggtcggatcctggtactggcaaaccgat<br>ggcatggaactgtacaaaggcggtcgtatgaataataccttctttcactcgaacgatgacgt<br>actgaaactttatcactcgaacgtaacggtagataacaccgtgatttggaaaaatgaaaatg<br>gtcctgttattcagtggggctgggccccgcacaacatcgacaacgtcgtggttaccaatacg<br>gacgtaattcataatcgcatgtattggaaagatgtgaagtataatacctgcattattaacag<br>ctcatcgcattacgcggatatgggcagcagcactactgcggatacggccaaaacaatttcga<br>acttccgtctggaaaacataacagttgaaggcatgacgaactgcgcagtgcgaatttatgcg<br>ttaagcaacaccgaaaacattcatatcaaaaatctgaatatcggcggttggaacggtcttga<br>ttacggctcccaagtctcgcatctgaaacgctactcggacacagccaacaacaaggtctggt<br>tgggcaacgaaactatcgatagtcggggcattaagctggagaattacaccgtgggcggcgtt<br>aaaatcgacaaaaccaccaccaattggaatgatacccagctcggtcggttaggctttgaccc<br>cgaaaattgggatacctggaacgcgtggtaactcgagcaaagcccgccgaaaggcgggcttt<br>tctgttgtttgagaccctactctggcgtcgatgaggga | 1960 |

Table S1 continued

| Construct | Sequence (5' → 3') | bp |
| --- | --- | --- |
| CeDex2365-49<br>J23119-UTR1-CeDex2365-49-T500 | caatccgccctcactacaaccgggtctctctatttgacagctagctcagtcctaggtataat<br>gctagcgctagcaataattttgtttaactttaagaaggagatataccatggcgccgcctgcg<br>accgtagcctccccggaagatggtgcggtcacactcgatcgcaccgcgaacacggatgattt<br>gcgcacatggtggcatcaggacacggcagcgaccacggatccgattgcggatgatcaagtac<br>gacagtctccgttctacgatgcaacgatcgccacgaccaccgatccgcaagatcagttcggc<br>tcgttcgtgtacttgtcggtgccacgctctggtaacgataaaattggctatacggaagaaga<br>tggagccgaattttcgagcgctgctgatctgaccatgagctggagctccttcgagtattcga<br>ccgatgtatgggtggatgttacgctgaaaaccggtcaggcaattacttcagccgatcaggtt<br>accatacggccgaccgatttaggcctacagacccagctagtcgatggtgacacggtgcgcgt<br>taaagtgccatttagcgaagaaggctatcgtttcagtgtggaattcgatccgcagcttatta<br>ccgcctacaatgacatgagtggtacgtcgggtaaattaaccacctcagccgaaggaaatcgt<br>gctattagcacggaaccgcaaaatagcatgctggtatttgcgaatcctaccccctccgcagc<br>ggaggctgagcgtacaatccccactgaacaatcaggcactatttaccgccctcagccgggac<br>tggttaacgacctggatgacatcacccaagaaattgtctactttggcccgggcacgtataac<br>atgggcgcggactaccacgctgtgctcccggagaatgtccgttgggtttatttagcaccagg<br>agcctatgtgaaaggcgcgtttcgcttccctcatgaaactcaaccacagttcaaagtgacgg<br>gtctgggcgtgctttctggagagcagtacgtctacgaagcggacaccagtaatggttatcgt<br>cacaacgttaatgacaactgtcacgtcacgtgcgtgaaaatgctgcaattcgaatccagcac<br>ccagccgcagacattggaactccagggggttactattaaccagccgccctatcattcttttg<br>tagtgtacgcacatgaaggggacaacgaggtgggcgtagagaattttcagatgaatgttgat<br>aactacaaacaggtgggcagctggtattggcagacagacggtattgaactgtacgacggcgg<br>ccgcatgaagaacacgttttttcacgcgaacgatgacgttctgaaaatgtatcacagcgatg<br>tcgcaatagaaaataccgttatctggaaaaatgaaaacggcccggtaatccagtggggctgg<br>actccgagggaaatccaagatgtgaccgtcaccaacaccgacgtgatccataatcgtatgta<br>ttggaaggatactaaatataacacctgtatttttaactcaagcagccactggcaggacatgg<br>gcgccaccgatcgagcggacacagcaaccactgtgaaaaatatgcgtttttcagacacccgc<br>gtggaaggcaccacaaactgcgcgcttcgcatttacgcgcttagtaataccgaaaacatcga<br>gattgacggtctgcatattgacgcgtggaacgagctggatgccgccgagcaggtcaatcagc<br>tgaagcgttatgcggatccggcaggtattaaggtgaccttaggcaatgaagttacggatggt<br>aatgggctgctgctgcataactatactgtaggaggtgttcctatcctgaaggcaggcgacaa<br>ttgggccagcactgaattggggcgcttagattttgatgccgatacatgggattcgtggaacg<br>ccaccagtgatgatcaacccgccgggccggcgtaactcgagcaaagcccgccgaaaggcggg<br>cttttctgttgtttgagaccctactctggcgtcgatgaggga | 2026 |

##### S1.3 Cell-free extract preparation

*E. coli* crude extract was prepared from BL21 Rosetta2 $\Delta recBCD$ (Addgene #176583) following the procedure of Silverman et al.,<sup>8</sup> with cell disruption performed using a Microfluidizer (LM10, Microfluidics) rather than bead beating. S30A and S30B buffers were prepared according to Sun et al.<sup>9</sup> The energy-buffer formulation was based on Sun et al., with exogenous tRNA omitted. Mg-glutamate and K-glutamate concentrations were adjusted to 4 and 110 mM, respectively, in the nominal final TXTL formulation. Cells were grown in 2×YTP medium at 37 °C and 200 rpm and harvested at an OD<sub>600</sub> of 2.0 ± 0.2. Cell pellets were washed three times with ice-cold S30A buffer and resuspended at 1 mL S30A per gram wet pellet. Cells were lysed by three passes through a microfluidizer at 800 bar. The lysate was supplemented with 4 mM DTT, clarified by centrifugation, subjected to an 80-min run-off reaction at 37 °C, and dialyzed against S30B buffer for 3 h at 4 °C using a 10-kDa MWCO dialysis cassette. Clarified extract aliquots were snap-frozen in liquid nitrogen and stored at −80 °C until use.

#### S1.4 Cell-free transcription–translation

Cell-free transcription–translation (TXTL) reactions were prepared using laboratory-made *E. coli* crude extract and energy buffer. Unless otherwise stated, formulations used for polymersome encapsulation contained 33% (v/v) crude extract, 33% (v/v) energy buffer, 5 nM linear mScarlet3 DNA, 70 mg mL<sup>-1</sup> Dex70, and nuclease-free water. Dextranase-containing reactions additionally contained 30 nM linear PsDex1711 DNA. FITC–mPEG 20 kDa was included at 10 μM where indicated. Reactions were prepared without pre-incubation and incubated at 28 °C.

#### S1.5 Bulk osmolality measurements

Bulk osmolality assays were used to establish that dextran depolymerization can generate an osmotic output under the reaction conditions used for the polymersome experiments and to select the dextranase construct and DNA concentration used in subsequent assays. All normalized comparisons use controls measured in the same experimental run. For the PsDex1711 DNA-concentration screen in Figure S4, $\Delta\Delta$ osmolality was calculated relative to the 0 nM PsDex1711 condition containing 5 nM mScarlet3 DNA, which therefore has a normalized value of zero by definition. The dextranase construct screen in Figure S3 was likewise normalized to the corresponding mScarlet3-only reference measured in the same run.

### S2 Microfluidic device, polymersome preparation, and imaging

#### S2.1 Device design and fabrication

The microfluidic device design is shown in Figure S1.

**SU-8 master mold fabrication.** Master molds were fabricated on 4-inch, single-side-polished (100) silicon wafers (525 μm nominal thickness; MicroChemicals, WSM40525250P1324XNN1) using SU-8 25 negative photoresist (Kayaku Advanced Materials) and maskless photolithography. The nominal channel-height target was approximately 20 μm, with master structures prepared in the 20–24 μm range. Wafers were cleaned with acetone and isopropanol, rinsed with deionized water, dried with nitrogen, and dehydrated at 200 °C for 5 min.

The photoresist was spin-coated using a LabSpin 6 BM spin coater (SÜSS). A spreading step at 500 rpm for 5 s with an acceleration of 100 rpm s<sup>-1</sup> was followed by spinning at 2000 rpm for 30 s with an acceleration of 300 rpm s<sup>-1</sup>. The coated wafers were soft-baked at 65 °C for 3 min and then at 95 °C for 10–15 min. The channel pattern was written using a μMLA maskless lithography system (Heidelberg Instruments) at an exposure dose of 210 mJ cm<sup>-2</sup>.

Post-exposure baking was performed at 65 °C for 1 min and 95 °C for 3 min. Unexposed resist was removed by immersion in propylene glycol monomethyl ether acetate (PGMEA; MicroChemicals, MPGU1050) for approximately 4 min. The developed masters were rinsed in isopropanol for 3 min and dried with nitrogen. The masters were then hard-baked at 150 °C for approximately 2 h, with the duration adjusted according to the structure geometry. Before PDMS casting, the masters were vapor-silanized under vacuum at room temperature for 2 h using 100 μL of methyltrichlorosilane (Sigma-Aldrich, 679208) to facilitate demolding.

**PDMS device fabrication.** PDMS devices and PDMS-coated glass substrates were prepared using a procedure adapted from Deshpande and Dekker.<sup>10</sup> PDMS base and curing agent (SYLGARD 184, Dow) were mixed at a 10:1 mass ratio and degassed in a vacuum desiccator. Approximately 30 g of the mixture was poured over each silanized SU-8 master, with an aluminum-foil rim used to contain the PDMS. After curing at 80 °C for 2 h, the PDMS replica was peeled from the master and cut into individual devices. Inlet and outlet holes were made using a 0.5-mm sampling punch (Micro to Nano, 52-004905).

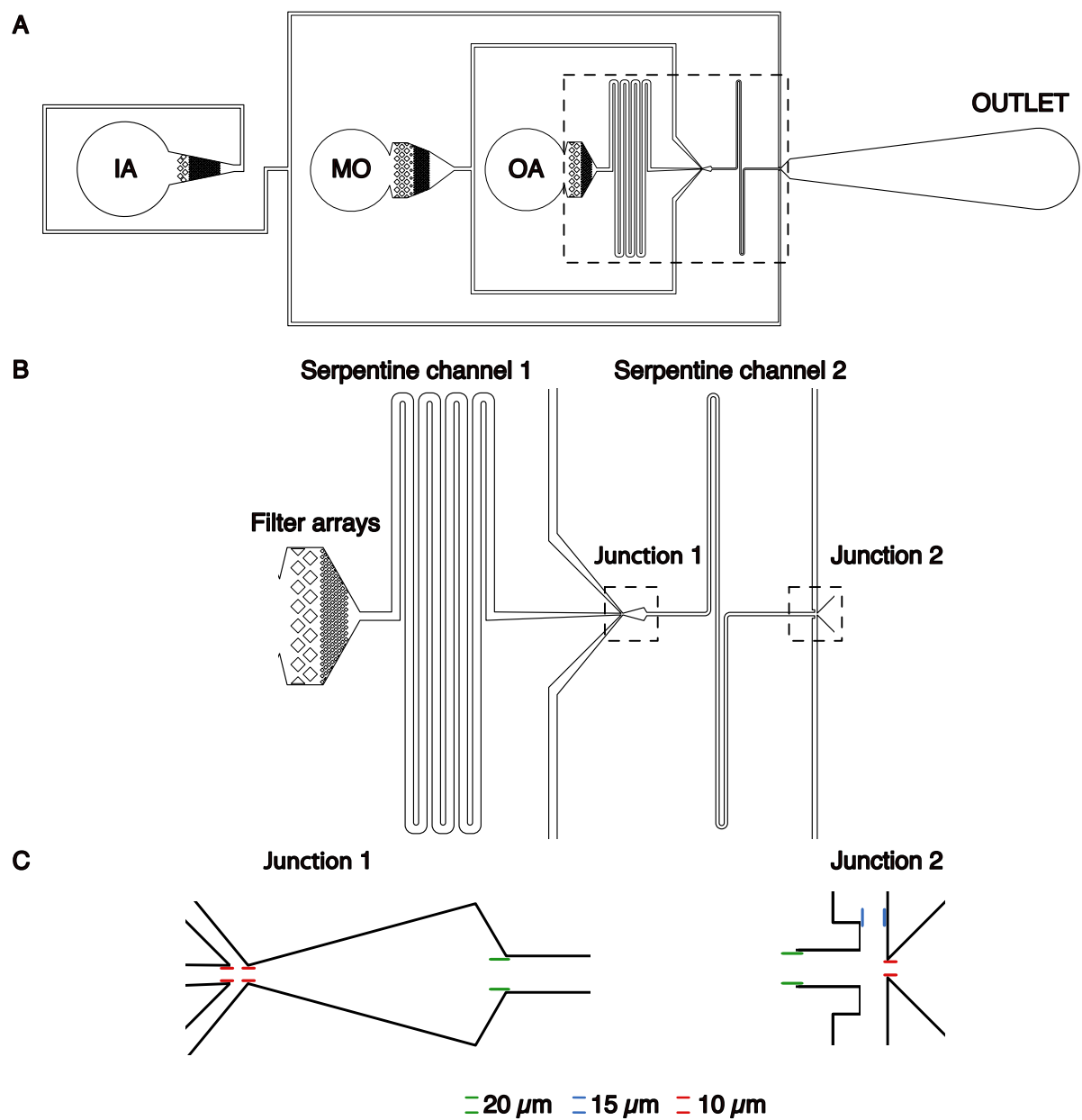

**Figure S1.** Microfluidic device design. (A) Complete CAD layout showing the inner aqueous (IA), middle oil (MO), and outer aqueous (OA) phase inlets and the outlet. (B) Enlarged view of the region outlined in (A), showing the filter arrays, serpentine channels, and two junctions. (C) Detailed views of the two junctions outlined in (B). Colored markers indicate the corresponding channel widths: green, 20 $\mu\text{m}$; blue, 15 $\mu\text{m}$; and red, 10 $\mu\text{m}$. Dimensions represent CAD design values.

**PDMS-coated glass substrates.** Two glass slides were coated using approximately 20 g of degassed PDMS mixture prepared at the same 10:1 mass ratio. A thin layer of PDMS was poured onto a silanized blank silicon wafer surrounded by an aluminum-foil rim. Glass slides were placed onto this layer and gently pressed to remove trapped air. The remaining PDMS was poured over the slides to fully cover them and form a level surface. After curing at 80 °C for 1 h, the PDMS-coated slides were released from the wafer using a blade.

**Device bonding.** The patterned PDMS surfaces and PDMS-coated glass substrates were activated by oxygen plasma (Nano, Diener electronic) for 40 s at a generator power setting of 40% and a process-pressure setpoint of 0.30 mbar. The activated surfaces were immediately brought into contact to bond. The assembled devices were heated at 120 °C for 10 min, followed promptly by selective PVA coating.

**Selective PVA surface treatment.** A 2% (w/v) aqueous PVA solution was introduced through the outer-phase inlet under microscopic observation. Air pressure of approximately 90 mbar was applied at the inner- and middle-phase inlets to prevent PVA solution from entering the upstream channels. After 4 min, the air pressure was increased sequentially to approximately 200 mbar at both inlets, and residual PVA solution was removed by air flushing and aspiration. The device was then baked at 120 °C for approximately 30 min.

#### S2.2 Phase compositions and polymersome preparation

**Solution preparation and handling.** A 20% (w/v) aqueous PVA stock solution was prepared by heating at 80 °C with stirring at 300 rpm overnight. The Dex1k stock was prepared by dissolving 1 g of Dex1k in 1 mL of water. Dex1k stock was added incrementally to adjust osmolality, with the number of adjustment and measurement cycles kept to a minimum. TXTL reaction mixtures were kept on ice until microfluidic encapsulation.

**Tracer-retention assay.** For the tracer-retention assay, the inner aqueous phase was PBS-based and contained 20 μM FITC-mPEG 2 kDa, 70 mg mL<sup>-1</sup> Dex70, and Dex1k for osmolality adjustment. The middle oil phase consisted of PL121 and 1-octanol (3:97, v/v). The outer aqueous phase was PBS-based and contained approximately 2% (w/v) PVA, with Dex1k added to adjust osmolality. The measured osmolalities of the inner and outer aqueous phases were 346 and 352 mOsm kg<sup>-1</sup>, respectively, giving an initial outer-minus-inner difference of 6 mOsm kg<sup>-1</sup>. No TXTL components were included.

#### S2.3 Image presentation

Microscopy recordings were stored in ND2 format. Fiji/ImageJ was used to inspect the recordings and select suitable display intensity ranges, which were then applied in Python during image preparation. Images for presentation were cropped using Python.

For the multi-polymersome images in Figure 2C of the main text, a square region encompassing the five selected polymersomes was used, with the same crop dimensions at both displayed time points. For the representative individual polymersomes in Figure 4A,B, crops were centered on the tracked particle positions. A common crop size was determined from the largest tracked radius across the selected experimental and control polymersomes plus a fixed margin, and applied across both sequences and all channels. The individual-polymersome crops were exported as 16-bit TIFF images without modifying pixel intensities.

For presentation, fixed minimum and maximum display intensities were applied within each channel throughout each displayed time series. Fluorescence images were pseudocolored cyan for FITC and magenta for mScarlet3. No background subtraction or gamma adjustment ($\gamma = 1$) was applied to the displayed microscopy images. Background correction was applied only during quantitative fluorescence analysis, as described in Section S3.

For the FITC-mPEG 2 kDa tracer-retention images in Figure 2A of the main text, the display ranges were 13604–65535 for brightfield and 2027–43648 for FITC. For Figure 2C of the main text, the display ranges were 15000–59000 for brightfield, 6000–52000 for FITC, and 1062–7000 for mScarlet3. For Figure 4A,B of the main text, the experimental sequence used display ranges of 1741–54497 for brightfield, 1011–15263 for FITC, and 617–2100 for mScarlet3, whereas the control sequence used 13791–56713 for brightfield, 6659–51172 for FITC, and 1245–6997 for mScarlet3.

For the supplementary microscopy sequences, the brightfield and mScarlet3 display ranges were 725–48800 and 650–1820 for EXP II (Figure S7), 2240–63100 and 710–2415 for EXP III (Figure S8), 730–51800 and 720–5710 for CTRL II (Figure S9), and 1208–66000 and 1010–6470 for CTRL III (Figure S10), respectively. For the expansion-with-retained-fluorescence example (Figure S11), the display ranges were 1586–42000 for brightfield and 650–1411 for mScarlet3.

#### S3 Image Analysis and Tracking Parameters

**Image Segmentation:** Image inputs for the Cellpose model combined contrast-limited adaptive histogram equalization (CLAHE)-enhanced brightfield images with normalized fluorescence images. To exclude artifacts and non-spherical structures, the resulting segmentation masks were filtered using strict geometric criteria, requiring a circularity $\geq 0.70$, a solidity $\geq 0.85$, and an eccentricity $\leq 0.85$. For each validated polymersome, the radius was calculated from the segmented contour perimeter $P$ as $r = \frac{P}{2\pi}$.

**Trajectory Reconstruction:** Polymersome trajectories were linked from the validated mask centroids using the trackpy library (0.7). The tracking algorithm was configured with a maximum search range of 120 pixels and a memory buffer of 7 frames to bridge transient detection losses and maintain continuous tracking across the time series. For Figure S5, only valid trajectories with the maximum number of recorded observations within each experiment were included.

**Fluorescence Quantification:** To correct for fluctuating background signals, an image-wide background baseline—defined as the 10th percentile of all pixel intensities in the respective channel—was dynamically subtracted from the mean internal intensity of each vesicle at every frame. The total background-corrected integrated fluorescence was subsequently calculated by multiplying this corrected mean intensity by the area of the corresponding polymersome mask.

**Software Environment:** Data processing and tracking were implemented using Python (3.13), relying on NumPy (2.4.4), Pandas (3.0.2), SciPy (1.17.1), scikit-image (0.26.0), and trackpy (0.7). Statistical analyses and plotting were performed using Python (3.13). The complete source code is publicly available via GitHub.<sup>11</sup>

#### S4 Supplementary Results

##### S4.1 Bulk Characterization

###### S4.1.1 Bulk TXTL Expression

Bulk TXTL expression was monitored using mScarlet3 fluorescence to confirm that the reaction remained transcriptionally and translationally active in the presence of PsDex1711 DNA. Representative fluorescence kinetics are shown in Figure S2.

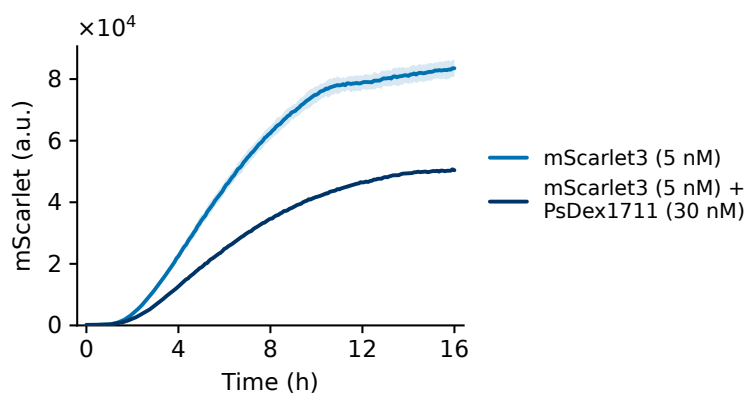

**Figure S2.** Bulk TXTL expression in the presence of PsDex1711 DNA. Representative mScarlet3 fluorescence kinetics from bulk TXTL reactions containing 5 nM mScarlet3 DNA either alone or together with 30 nM PsDex1711 DNA. Reactions were incubated at 28 °C. Lines show the mean fluorescence and shaded regions indicate the standard deviation of three technical replicates. Addition of PsDex1711 DNA reduced mScarlet3 reporter output but did not abolish TXTL activity.

###### S4.1.2 Dextranase construct screen

Three dextranase constructs were compared in bulk TXTL reactions to identify a construct capable of generating a measurable osmotic response in the presence of Dex70. The screen included the dextranase from *Arthrobacter* sp. CB-8, PsDex1711, and CeDex2365-49. Osmolality changes were normalized to the corresponding mScarlet3-only reference measured in the same experiment. Based on this screen, PsDex1711 was selected for subsequent polymersome experiments (Figure S3).

###### S4.1.3 Bulk osmolality responses

Additional bulk osmolality responses are shown in Figure S4.

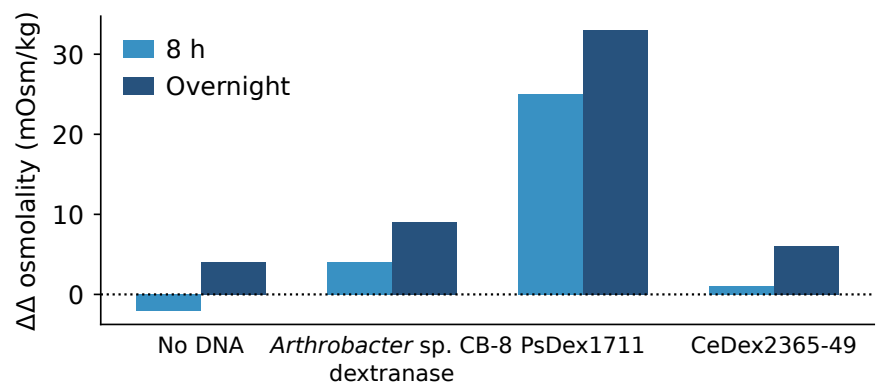

**Figure S3.** Bulk osmolality screen of dextranase constructs. Bulk TXTL reactions containing 70 mg mL<sup>-1</sup> Dex70 and 5 nM mScarlet3 DNA were supplemented with 20 nM DNA encoding the *Arthrobacter* sp. CB-8 dextranase, PsDex1711, or CeDex2365-49. The additional change in osmolality ($\Delta\Delta$) after 8 h and overnight incubation is shown relative to the corresponding same-run mScarlet3-only reference. The mScarlet3-only reference is omitted because its normalized value is zero by definition; the no-DNA condition is shown separately. PsDex1711 was selected for subsequent polymersome experiments. Each condition was measured once in a single experiment.

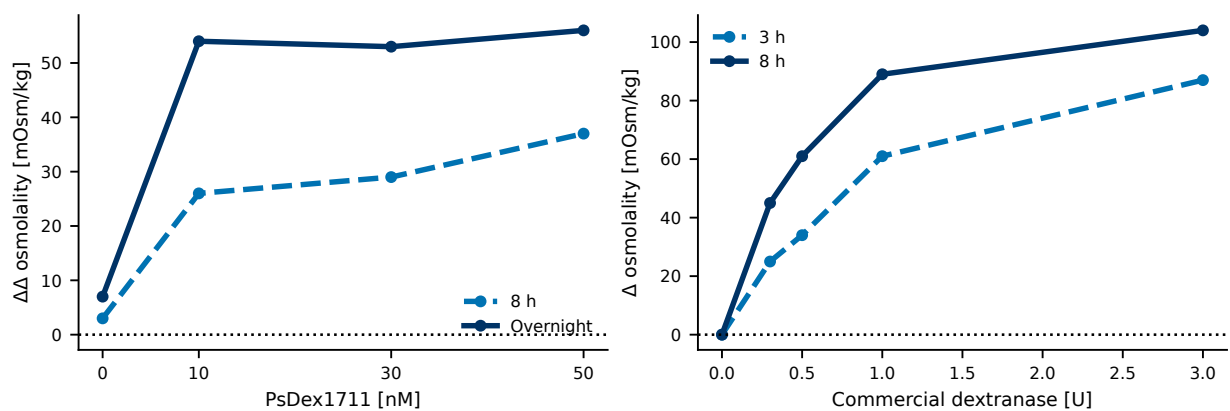

**Figure S4.** Dose-dependent osmotic responses generated by PsDex1711 expression and commercial dextranase. Left: PsDex1711 DNA-concentration screen in bulk TXTL reactions. The additional change in osmolality ($\Delta\Delta$) at 8 h and overnight is shown relative to the same-run 0 nM PsDex1711 reference containing 5 nM mScarlet3 DNA. PsDex1711 concentrations of 10, 30, and 50 nM were tested to identify a concentration producing a measurable osmotic output while retaining the mScarlet3 reporter in the formulation. Right: Bulk reactions containing dextran 70 kDa were incubated with increasing amounts of commercial dextranase. Commercial dextranase reactions were prepared in water without added buffer. The change in osmolality from the initial measurement ($\Delta$ osmolality) is shown after 3 and 8 h. The 0 U condition is the negative control measured in the same experiment. Each panel shows data from a single experiment.

#### S4.2 Population expansion across independent experiments

Two additional independent experiments per condition, performed on separate days, reproduced the direction of the population-level response shown in Figure 3C of the main text (Figure S5; Table S2). In the supplementary experiments, the PsDex1711-negative populations contracted, reaching endpoint mean normalized radii of 0.928 and 0.917. The PsDex1711-positive populations initially contracted and subsequently re-expanded to endpoint mean normalized radii of 1.006 and 1.010. Across all three independent experiments per condition, endpoint mean normalized radii were below unity without PsDex1711 and above unity with PsDex1711, although the magnitude of net expansion varied between experiments. Across all nine pairwise comparisons of individual-polymersome endpoint radius distributions between the three PsDex1711-negative and three PsDex1711-positive experiments, values were significantly higher in the presence of PsDex1711 (all $p < 0.001$, Mann–Whitney U tests).

**Table S2.** Population statistics for the main-text experiment (Figure 3C) and two additional independent experiments per condition (Figure S5). Values are means ± standard deviations. Minimum normalized radii summarize the individual-polymersome minima within the analysis window; times in parentheses are the mean times at which those minima occurred. Endpoint times were 14 h for the main-text experiment, approximately 14.5 h for SI A, and approximately 12.3 h for SI B.

| Condition | Date | $n$ particles | $n$ videos | Minimum normalized radius (time) | Endpoint normalized radius |
| --- | --- | --- | --- | --- | --- |
| +PsDex1711 (main) | 2026-07-03 | 145 | 38 | $0.978 \pm 0.028$ (2.7 h) | $1.055 \pm 0.067$ |
| +PsDex1711 (SI A) | 2026-06-05 | 847 | 29 | $0.988 \pm 0.011$ (3.9 h) | $1.006 \pm 0.025$ |
| +PsDex1711 (SI B) | 2026-07-05 | 639 | 54 | $0.989 \pm 0.014$ (2.8 h) | $1.010 \pm 0.026$ |
| –PsDex1711 (main) | 2026-07-08 | 150 | 32 | $0.931 \pm 0.048$ (11.8 h) | $0.933 \pm 0.048$ |
| –PsDex1711 (SI A) | 2026-05-09 | 245 | 16 | $0.925 \pm 0.028$ (13.6 h) | $0.928 \pm 0.031$ |
| –PsDex1711 (SI B) | 2026-04-02 | 31 | 19 | $0.916 \pm 0.021$ (11.9 h) | $0.917 \pm 0.022$ |

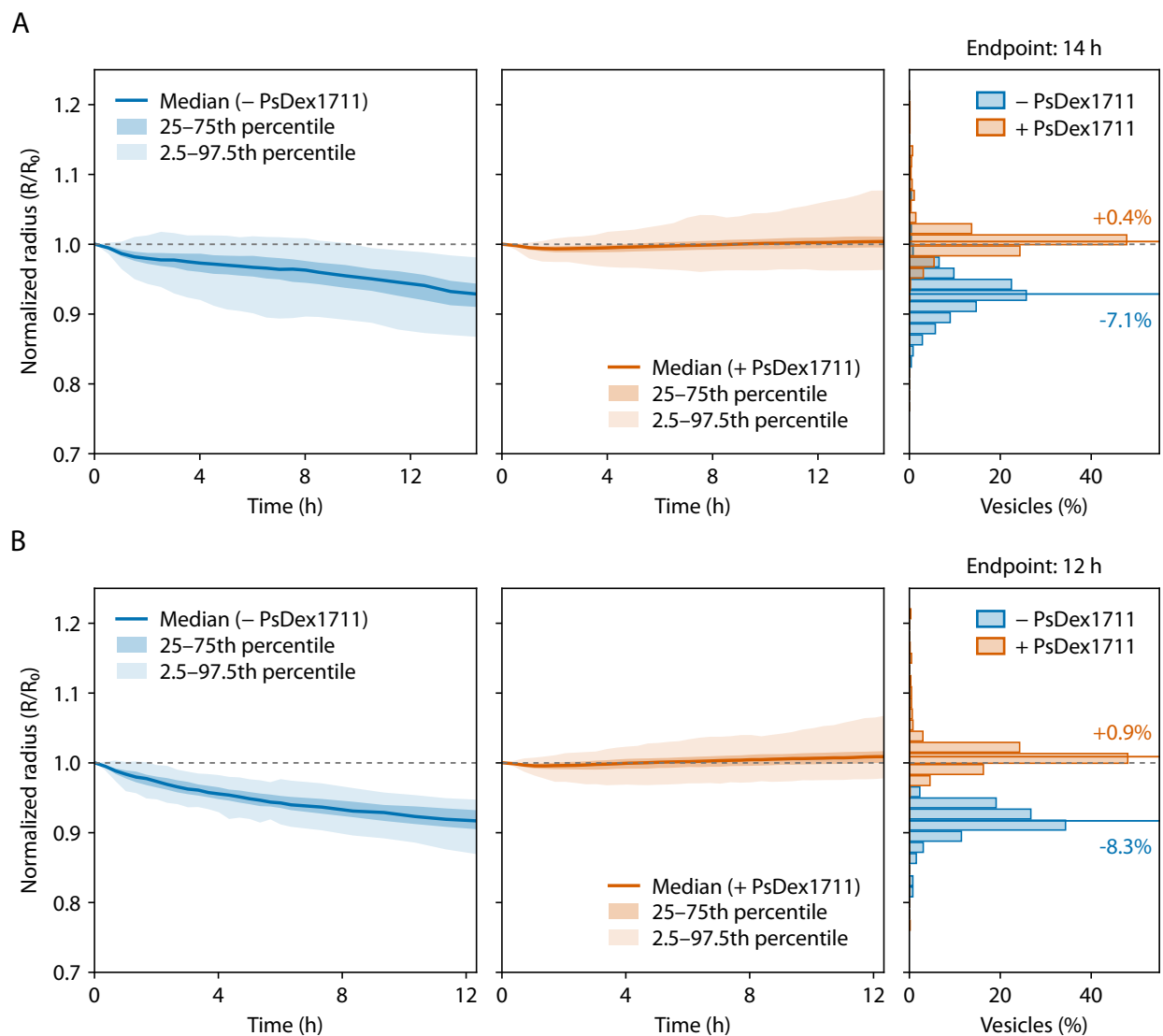

**Figure S5.** Population radius responses in two additional independent experiments per condition. Each row shows normalized radius trajectories ($R/R_0$) for mScarlet3-only controls without PsDex1711 DNA (left, blue), PsDex1711-containing polymersomes (middle, orange), and the corresponding endpoint distributions (right). Solid curves show medians; darker and lighter shading indicate the 25th–75th and 2.5th–97.5th percentile ranges, respectively. Dashed horizontal lines mark $R/R_0 = 1$. Horizontal colored lines in the endpoint distributions mark medians, and percentages indicate median changes relative to the initial radius. (A) Control experiment dated 2026-05-09 and PsDex1711 experiment dated 2026-06-05, evaluated at approximately 14.5 h. (B) Control experiment dated 2026-04-02 and PsDex1711 experiment dated 2026-07-05, evaluated at approximately 12.3 h. Plot headings use integer-hour labels. Sample sizes and mean radius statistics are given in Table S2.

##### S4.3 Distribution of peak-normalized fluorescence

To assess whether polymersomes lost internally expressed mScarlet3 without completely rupturing, we compared the integrated fluorescence of each intact polymersome near the end of the experiment (12.3–13 h) to its individual peak value ($F_{\text{end}}/F_{\text{peak}}$). For PsDex1711-containing polymersomes ($n = 145$, evaluated at 13 h), the distribution revealed two distinct groups. The vast majority (97%) formed a main population that retained a median of 59% of its peak fluorescence. However, a small subset of five polymersomes (3%) exhibited a pronounced fluorescence loss, retaining at most 22% (median 14%) of their peak intensity. This subset is clearly separated from the main population by a gap and shows pronounced loss of internal fluorescence, consistent with possible content release (see Figure 4 in the main text for representative examples).

In contrast, the negative control ($n = 31$, evaluated at 12.3 h) exhibited a single, uniform population. These polymersomes retained a median endpoint fluorescence of 89% of the individual peak value, with no individual polymersome dropping below 77% of its maximum intensity. No comparably low-fluorescence subset was observed among the analyzed control polymersomes. Note that completely ruptured polymersomes in both conditions collapsed to sizes below the segmentation threshold and were naturally excluded from this analysis.

In Figure 4 of the main text, we show individual polymersomes from the negative-control population in Figure S6B to contrast them with the PsDex1711-containing polymersomes that exhibited fluorescence loss.

##### S4.4 Additional single-polymersome observations

###### S4.4.1 Additional examples of expansion-associated fluorescence loss

Additional image sequences and trajectories are shown for the EXP II, EXP III, CTRL II, and CTRL III polymersomes plotted in Figure 4 of the main text (Figures S7–S10). EXP II and EXP III exhibited maximum radius increases of approximately 11% and 18%, respectively, followed by pronounced mScarlet3 fluorescence loss. At the late image time points, approximately 9–10 h, background-corrected total fluorescence had decreased to approximately 22% and 18% of the respective peak values. In contrast, CTRL II and CTRL III contracted while retaining approximately 91% and 90% of their peak fluorescence at approximately 9 h.

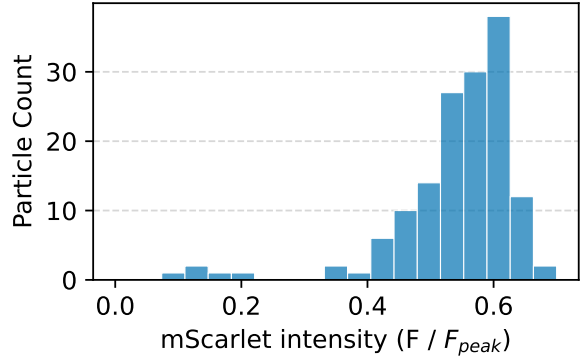

(a) PsDex1711-containing polymersomes.

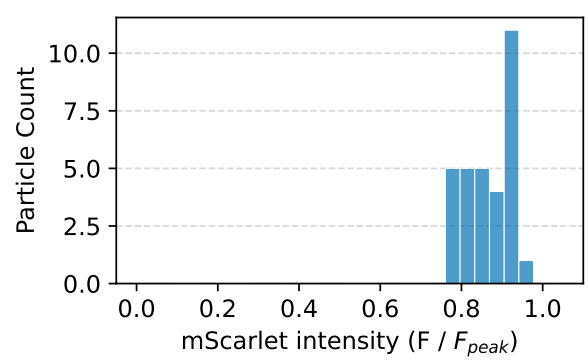

(b) Negative control.

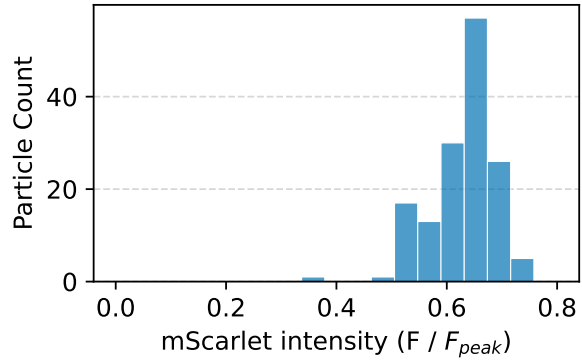

(c) Negative control from Figure 3C of the main text.

**Figure S6.** Distribution of peak-normalized endpoint fluorescence. (A) Histogram of background-corrected, integrated mScarlet3 fluorescence at 13 h, normalized to each polymersome's own peak value ($F/F_{peak}$), for $n = 145$ PsDex1711-containing polymersomes. A small, distinct subset of polymersomes is separated from the main population, indicating pronounced loss of internal mScarlet3 fluorescence. (B) Corresponding histogram for the negative control ($n = 31$, evaluated at 12.3 h; experiment dated 2026-04-02, as in Figure S5B). The distribution shows a single population with high fluorescence retention, with no comparably low-fluorescence subset in this analyzed sample. (C) Corresponding histogram for the negative-control population shown in Figure 3C of the main text ($n = 150$, evaluated at 13 h). For all data, fluorescence was integrated within each polymersome mask after subtracting a full-image background estimate.

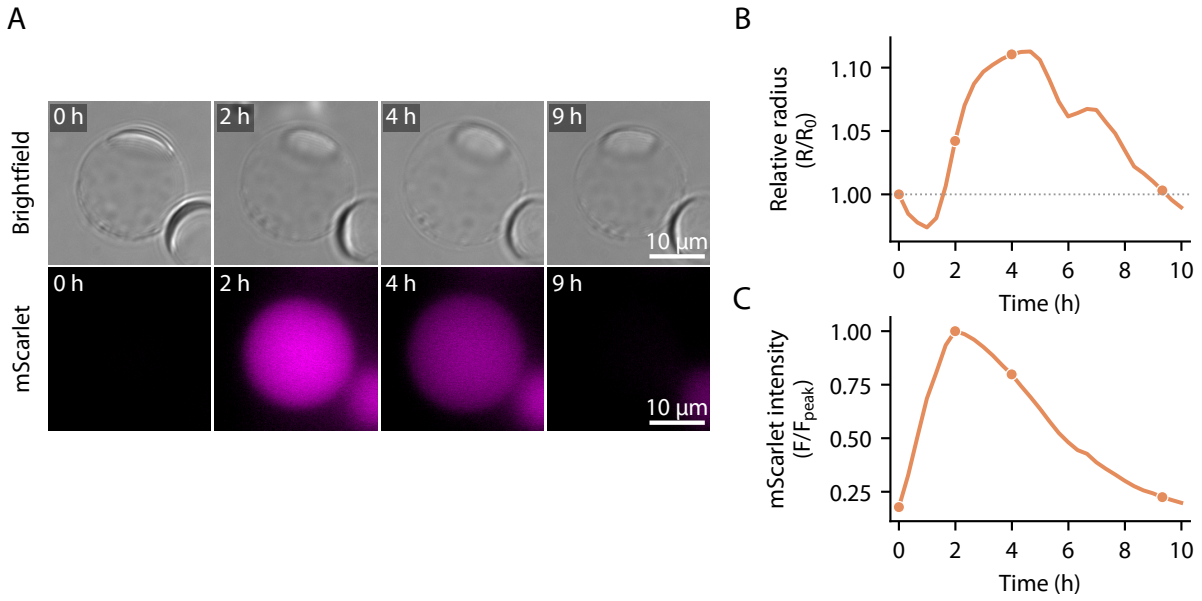

**Figure S7.** Expansion and fluorescence loss in EXP II, a selected PsDex1711-containing polymersome plotted in Figure 4 of the main text. (A) Brightfield and mScarlet3 fluorescence images at approximately 0, 2, 4, and 9 h. Image labels indicate approximate elapsed times. Fluorescence is shown in magenta. Scale bars, 10 $\mu\text{m}$. (B) Radius normalized to its initial value ($R/R_0$); the dotted line marks $R/R_0 = 1$. (C) Background-corrected total mScarlet3 fluorescence normalized to its maximum over the full retained time series ($F/F_{\text{peak}}$). Circular markers in (B,C) identify the frames displayed in (A). All panels describe the same polymersome.

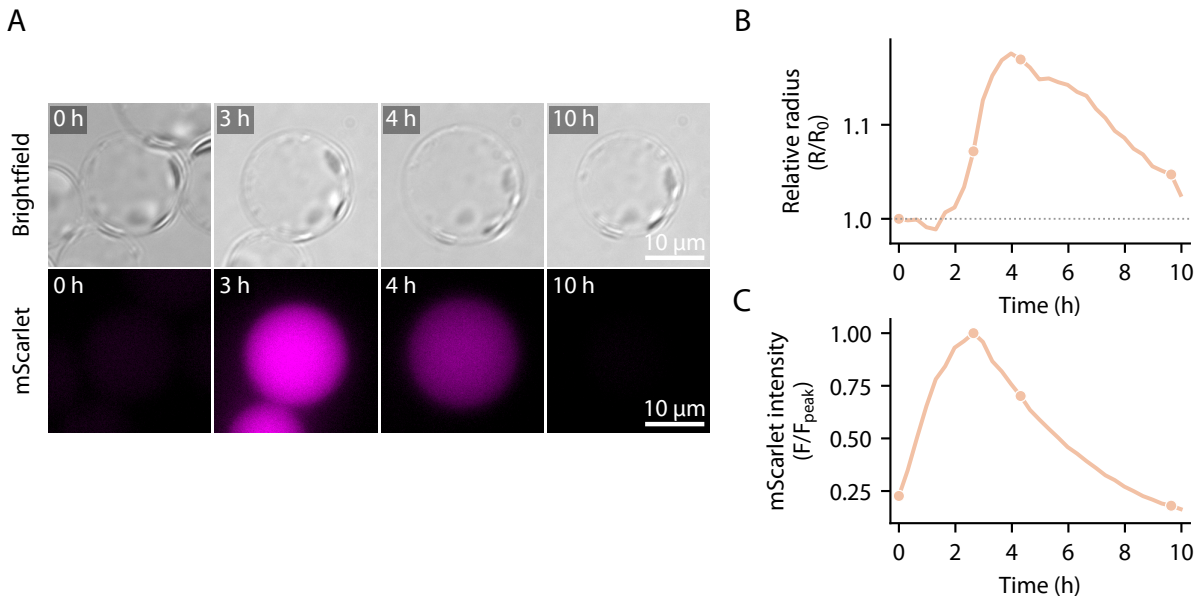

**Figure S8.** Expansion and fluorescence loss in EXP III, a selected PsDex1711-containing polymersome plotted in Figure 4 of the main text. (A) Brightfield and mScarlet3 fluorescence images at approximately 0, 3, 4, and 10 h. Image labels indicate approximate elapsed times. Fluorescence is shown in magenta. Scale bars, 10 $\mu\text{m}$. (B) Radius normalized to its initial value ($R/R_0$); the dotted line marks $R/R_0 = 1$. (C) Background-corrected total mScarlet3 fluorescence normalized to its maximum over the full retained time series ($F/F_{\text{peak}}$). Circular markers in (B,C) identify the frames displayed in (A). All panels describe the same polymersome.

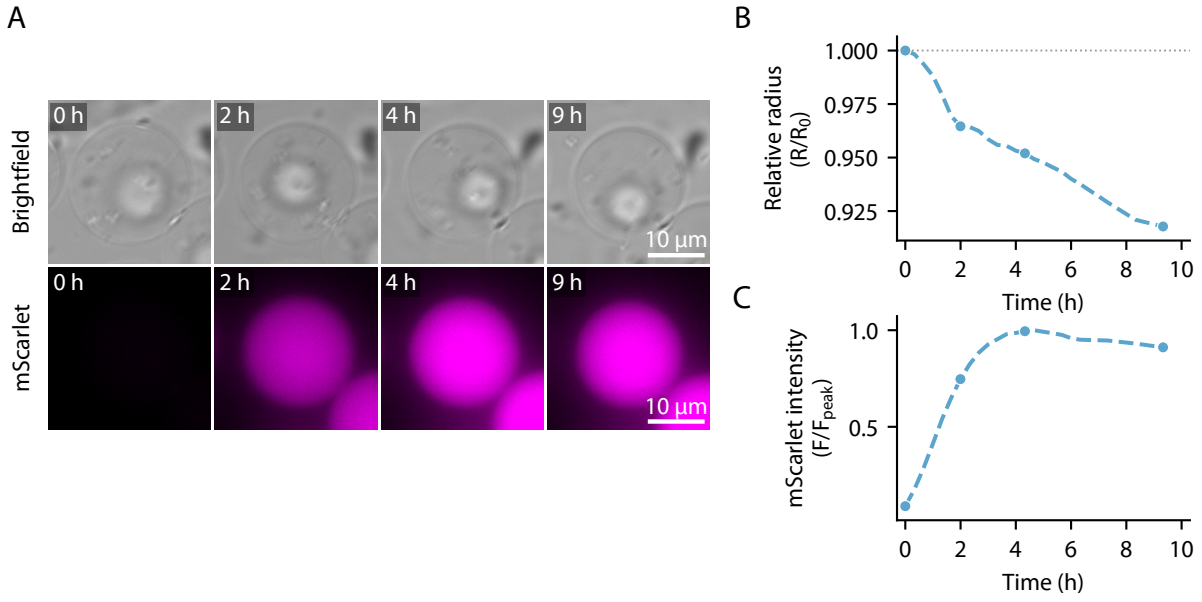

**Figure S9.** Contraction with retained fluorescence in CTRL II, a selected mScarlet3-only control polymersome without PsDex1711 DNA, plotted in Figure 4 of the main text. (A) Brightfield and mScarlet3 fluorescence images at approximately 0, 2, 4, and 9 h. Image labels indicate approximate elapsed times. Fluorescence is shown in magenta. Scale bars, 10 $\mu\text{m}$. (B) Radius normalized to its initial value ($R/R_0$); the dotted line marks $R/R_0 = 1$. (C) Background-corrected total mScarlet3 fluorescence normalized to its maximum over the full retained time series ($F/F_{\text{peak}}$). Circular markers in (B,C) identify the frames displayed in (A). All panels describe the same polymersome.

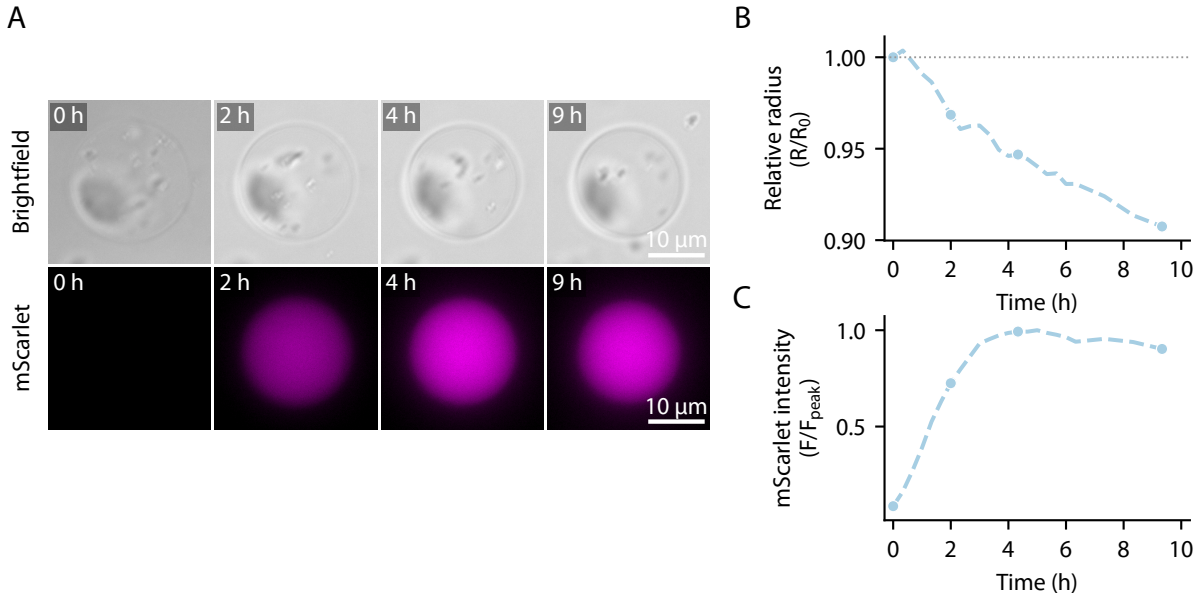

**Figure S10.** Contraction with retained fluorescence in CTRL III, a selected mScarlet3-only control polymersome without PsDex1711 DNA, plotted in Figure 4 of the main text. (A) Brightfield and mScarlet3 fluorescence images at approximately 0, 2, 4, and 9 h. Image labels indicate approximate elapsed times. Fluorescence is shown in magenta. Scale bars, 10 $\mu\text{m}$. (B) Radius normalized to its initial value ($R/R_0$); the dotted line marks $R/R_0 = 1$. (C) Background-corrected total mScarlet3 fluorescence normalized to its maximum over the full retained time series ($F/F_{\text{peak}}$). Circular markers in (B,C) identify the frames displayed in (A). All panels describe the same polymersome.

###### S4.4.2 Expansion with retained fluorescence

A selected PsDex1711-containing polymersome exhibited pronounced expansion while retaining strong internal mScarlet3 fluorescence (Figure S11). Its radius increased by approximately 25% relative to its initial value, while integrated fluorescence remained at approximately 93% of its peak at 10 h. This example shows that substantial expansion was not necessarily accompanied by pronounced fluorescence loss.

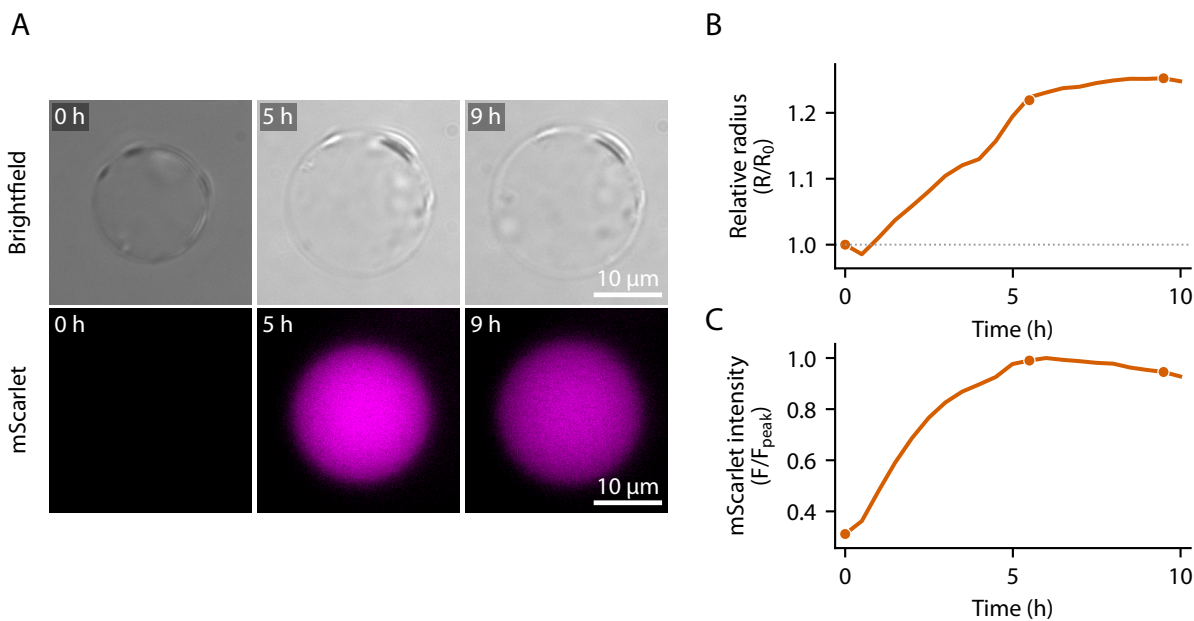

**Figure S11.** Pronounced polymersome expansion with retained mScarlet3 fluorescence. (A) Brightfield and mScarlet3 fluorescence images of a selected PsDex1711-containing polymersome at approximately 0, 5, and 9 h. Image labels indicate approximate elapsed times. Fluorescence is shown in magenta. Scale bars, 10 $\mu\text{m}$. (B) Radius normalized to its initial value ($R/R_0$); the dotted line marks $R/R_0 = 1$. (C) Background-corrected integrated mScarlet3 fluorescence normalized to its maximum over the recorded time series ($F/F_{\text{peak}}$). All panels describe the same polymersome.

###### S4.5 FITC–mPEG fluorescence under different imaging schedules

To assess the contribution of photobleaching to the fluorescence decline discussed in Figure 2A,B of the main text, we compared FITC–mPEG fluorescence under two imaging schedules (Figure S12). The frequently imaged polymersome showed a greater relative fluorescence decrease, retaining approximately 75% of its initial signal at the final recorded time point, compared with approximately 88% under sparse imaging. This observation is consistent with a contribution from photobleaching during repeated image acquisition.

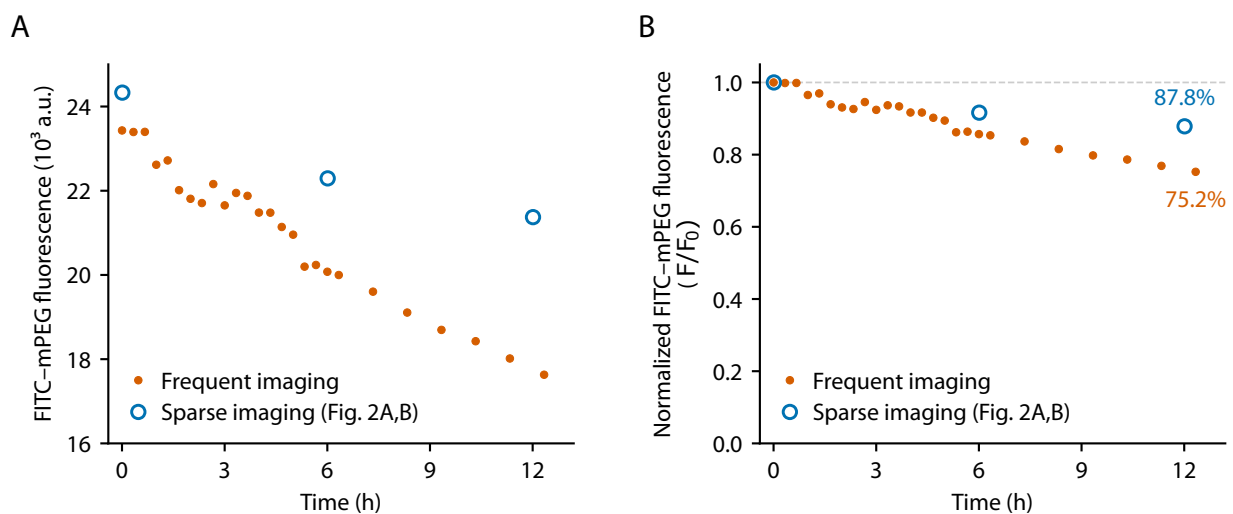

**Figure S12.** FITC-mPEG fluorescence under different imaging schedules. (A) Absolute fluorescence and (B) the same data normalized to the initial fluorescence of each series ($F/F_0$). Each series follows a single polymersome selected from an isolated region without neighboring polymersomes. The frequent-imaging series (orange filled circles) was acquired every 20 min during the first 6 h and every 1 h thereafter. The sparse-imaging series (blue open circles) corresponds to the polymersome shown in Figure 2A,B of the main text, imaged at 0, 6, and 12 h. Fluorescence declined more rapidly under frequent imaging, consistent with a contribution from photobleaching. Percentages in (B) indicate the remaining normalized fluorescence at the final recorded time point of each series; the dashed line marks $F/F_0 = 1$.

#### Supplementary Tables

**Table S3.** Commercial reagents and materials used in this study.

| Product | Manufacturer / supplier | Catalog number / grade |
| --- | --- | --- |
| Pluronic L121 | Sigma-Aldrich | 435457 |
| 1-Octanol | Sigma-Aldrich | 297887 |
| Poly(vinyl alcohol), Mowiol 4-88 | Sigma-Aldrich | 81381 |
| Dextran 1 kDa | MedChemExpress | HY-112624A |
| Dextran T70 (70 kDa) | MedChemExpress | HY-112624B |
| Ficoll PM70 | GE Healthcare | 17-0310-10 |
| FITC-mPEG 2 kDa | MedChemExpress | HY-D2840D |
| FITC-mPEG 20 kDa | MedChemExpress | HY-D28401 |
| PBS buffer, 10×, pH 7.4 | Invitrogen (Thermo Fisher Scientific) | AM9625 |
| Anti-Evaporation Oil | ibidi | 50051 |
| PDMS silicone elastomer kit | Dow | SYLGARD 184 |
| repliQa HiFi ToughMix | Quantabio | 95200-500 |
| NucleoSpin Gel and PCR Clean-up | Macherey–Nagel | 740609.250 |
| BL21 Rosetta2 $\Delta recBCD$ | Addgene | #176583 |
| Sampling punch, 0.5 mm | Micro to Nano | 52-004905 |
| Methyltrichlorosilane | Sigma-Aldrich | 679208 |
| Silicon wafer, 4 inch, (100), single-side polished | MicroChemicals | WSM40525250P132<br>4XNN1 |
| PGMEA | MicroChemicals | MPGU1050 |
| Dextranase from *Penicillium* sp. | Sigma-Aldrich | D4668-1KU |
